# Physical confinement selectively favours bacterial growth based on cell shape

**DOI:** 10.1101/2024.05.06.592621

**Authors:** M Sreepadmanabh, Meenakshi Ganesh, Pratibha Sanjenbam, Christina Kurzthaler, Deepa Agashe, Tapomoy Bhattacharjee

## Abstract

How are bacterial communities altered by changes in their microenvironment? Evidence from homogeneous liquid or flat plate cultures implicates biochemical cues — such as variation in nutrient composition ^1,2^, response to chemoattractants and toxins ^3,4^, and inter-species signalling ^5,6^ — as the primary modes of bacterial interaction with their microenvironment. However, these systems fail to capture the effect of physical confinement on bacteria in their natural habitats. Bacterial niches like the pores of soil, mucus, and infected tissues are disordered microenvironments with material properties defined by their internal pore sizes and shear moduli^7–11^. Here, using three-dimensional matrices that match the viscoelastic properties of gut mucus, we test how altering the physical properties of their microenvironment influences bacterial growth under confinement. We find that low aspect-ratio bacteria form compact, spherical colonies under confinement while high aspect-ratio bacteria push their progenies further outwards to create elongated colonies with a higher surface area, enabling increased access to nutrients. As a result, the population level growth of high aspect-ratio bacteria is more robust to increased physical confinement compared to that of low aspect-ratio bacteria. Thus, our results capture experimental evidence showing that physical constraints can play a selective role in bacterial growth based on cell shape.

## 1. Introduction

Most natural habitats host diverse bacterial communities, which actively respond to extrinsic environmental cues that reflect the dynamic properties of their microenvironments ^7,12^. Identifying the basis of these feedback-response cycles is critical for describing the evolution of natural microbial populations, characterizing host-symbiont crosstalk and interdependency, as well as understanding pathogenesis to design effective antibiotics. A major class of such interactions is understood within the framework of biochemical signalling, where small molecules (such as metabolites, growth factors, antibiotics, and chemoattractants) drive cellular perception and consequent responses ^1–4^. These processes also directly impact the composition of natural microbial communities, wherein co-existing species exhibit symbiotic, mutualistic, or predatory interactions with each other ^5,6^. However, current experimental approaches designed to explore such interactions rely on bacteria cultured in homogeneous liquid broths or on surfaces of 2D flat plates, which do not capture the structural complexity of many natural niches. Bacteria often reside in complex and disordered 3D microenvironments like soil, inter-tissue pores, and biological hydrogels such as mucus, which feature a wide range of material properties ^9,13,14^. For example, the porosity and stiffness of soil is affected by its moisture content and granularity ^15^. The spatial architecture of tissues also undergoes significant structural alterations as a result of ECM (extracellular matrix) deposition and enzymatic remodelling by fibroblasts and immune cells^16^. Similarly, the viscoelastic properties of mucus vary with diet, inflammatory responses, microbial enzymatic activity, and pathologies like cystic fibrosis ^8,17,18^ (**Fig. 1a**). Furthermore, bacterial colonies physically constrained within these complex 3D matrices experience diffusion-limited growth ^19,20^. In contrast, bacteria growing in homogeneously well-mixed liquid cultures have unrestricted access to nutrients, while colonies on a flat agar surface receive oxygen supply from the top as well as direct nutrient access from the underlying substratum. Although diffusion-limited zones can be established during growth over time, these usually affect the colony’s core once multilayer 3D growth occurs, with cells piling on top of each other ^21^. However, growth under 3D confinement could introduce such consumption-diffusion limited zones locally around small single cell-derived colonies, despite there being a global abundance of the nutrients in the bulk medium. Hence, growth under 3D confinement becomes fundamentally different from that on standard 2D interfaces. Previous work has investigated colony growth under nutrient-replete 2D conditions, extending growth-generated stresses as a putative mechanism for achieving quasi-3D conformations ^22,23^. However, these studies do not fully capture bacterial growth in disordered, confined microenvironments. Growing evidence indicates that bacterial motility is significantly altered by confinement within a granular and porous matrix ^24^ and the growth of initially smooth, densely-packed bacterial populations under 3D confinement shows generic surface roughening morphodynamics driven by differential nutrient accessibility^20^. Despite this, bacterial population level growth and colony organization under complex and disordered confinement remains insufficiently described. Consequently, whether and how the physical constraints imposed by a 3D growth microenvironment influence microbial populations remains unclear.

**Figure 1:**
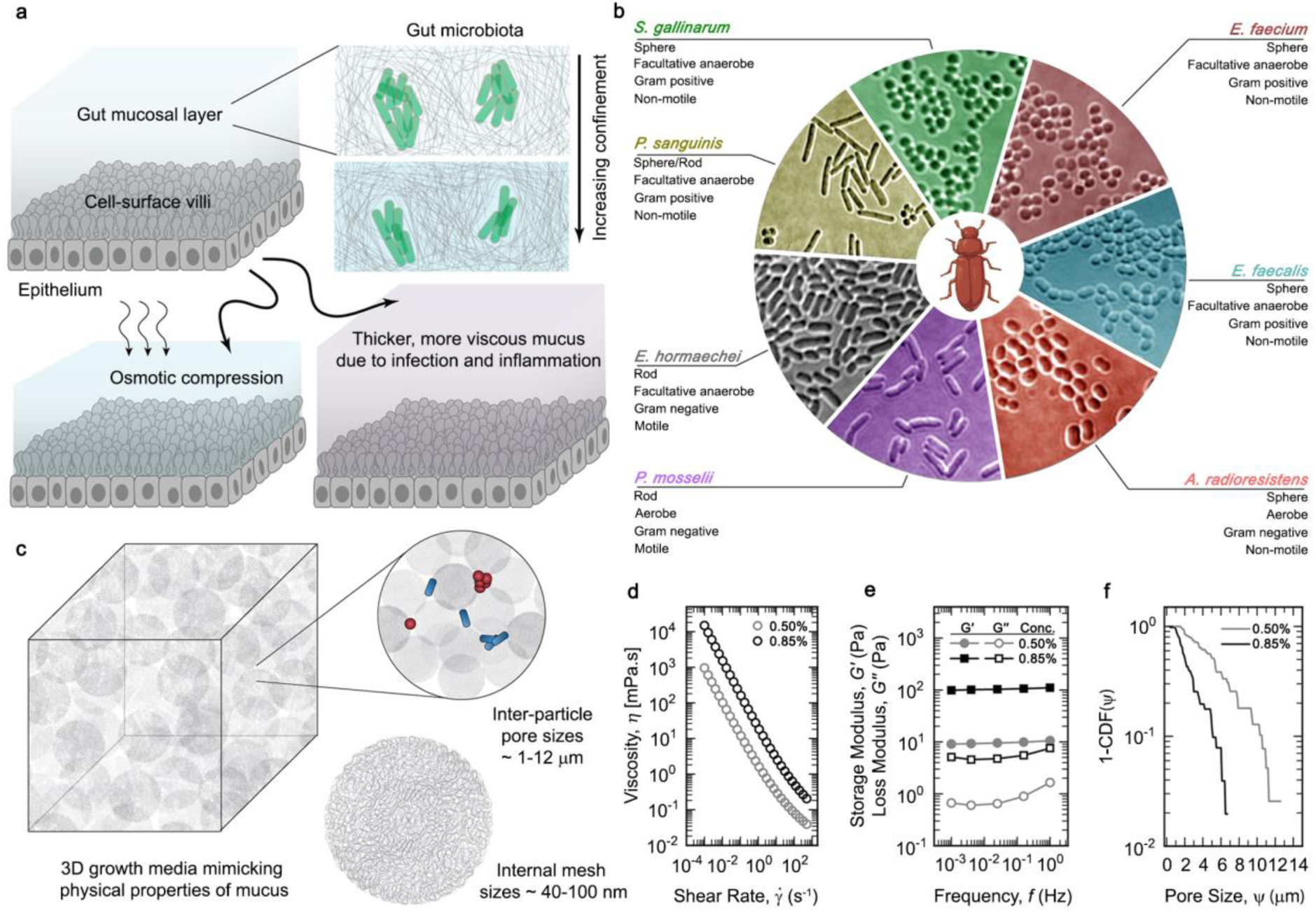
An *in vitro* 3D growth matrix with tunable viscoelastic properties. (a) The physical properties of the mucosal layer are diverse and are altered by factors such as a change in diet, infection, inflammation, and enzymatic degradation of fibers by microbes. (b) Different gut-derived microbial strains from red flour beetles, imaged in brightfield (pseudo-colored for enhanced contrast). (c) By packing highly swollen polymeric hydrogel granules beyond jamming concentrations, we design an *in vitro* 3D growth medium, which provides a granular and internally porous microenvironment for bacterial growth under confinement. (d-f) The physical properties of the microgel growth medium are highly tunable based on the mass percentage of hydrogel granules relative to liquid LB (%weight/volume). Here, we show both soft and low confinement (0.50%) as well as stiff and high confinement (0.85%) matrices, the viscoelastic properties of which approximately match mucosal samples from natural sources ^26^. Rheological measurements shown here either apply (d) a unidirectional shear at different rates to measure the viscosity, or, (e) a small amplitude (1%) of oscillatory strain at different frequencies to measure the shear moduli of the 3D growth media. The storage modulus (*G’*) is a measure of the elastic solid-like nature, while the loss modulus (*G’’*) signifies the viscous properties of the material. (f) The porosity of the medium, shown here as a complementary cumulative distribution function (1-CDF) of all the inter-particle pore spaces, determines the degree of confinement, and can be tuned by altering the packing fraction of the hydrogel granules.

In this work, we present experimental evidence for confinement-dependent growth dynamics that selectively favour specific bacterial morphologies. We prepare transparent, granular 3D growth media—a special type of porous matrix for direct measurement of bacterial growth as well as visualization of colony organisation under different biomimetic degrees of confinement—that broadly mimic the structural and viscoelastic properties of mucus, a biological hydrogel. Using several different bacterial strains isolated from the gut mucus of red flour beetles we show how an increase in confinement confers a growth advantage to bacteria with a higher aspect ratio. Our study combines quantitative measurements of bacterial growth and colony morphology, agent-based modelling, numerical calculations, as well as *in vitro* co-culture experiments, to establish a generalized principle for growth under confinement. By experimentally manipulating cellular morphology, we also demonstrate a remarkable interconvertible behaviour between high and low aspect ratio forms of the same bacterial species, which strongly implicates single-cell morphology as a broad determinant of colony architecture and growth success under physical confinement. Importantly, we show that population-level variation in growth dynamics can arise without invoking mechanisms such as genetic mutations, behavioural differences, or cellular responses to biochemical cues. Rather, our work suggest that the composition of heterogeneous microbial populations may be strongly dependent on efficient colony organization under increased confinement that selectively favours bacterial species with a high aspect ratio morphology. These principles are valuable for the experimental elucidation and theoretical modelling of microbial dynamics in complex, spatiotemporally varying natural niches.

## 2. Results

### 2.1 Mimicking physical properties of the mucosal microenvironment

To understand how the physical properties of their microenvironment regulate bacterial growth, we avoid the use of laboratory strains to minimize potential experimental artefacts arising from laboratory adaptation, such as mutations accumulated during long-term culture in homogeneous liquid cultures or 2D surfaces. Hence, we isolate several bacterial strains from the gut of the red flour beetle *(Tribolium castaneum*) and identify them using 16S rRNA sequencing. These bacteria naturally reside in a viscoelastic microenvironment, which ensures minimal interference from culturing practices. Together, these strains include a diverse representation of different envelope properties, metabolic modes, motility patterns, and morphologies (**Fig. 1b**). To create a structural mimic of their natural habitat, we design a jammed microgel-based 3D growth medium that mimics the physical and structural properties of mucus (**Fig. 1c**). Previous structural analysis of mucus indicates different regimes of porosities, spanning from nanometer-scale to micron-scale pores ^25^. Further, rheological measurements on mucosal samples from various sources also reveal a broad range of shear moduli (ranging from ∼ 1 Pa to ∼ 100 Pa) and viscosities (ranging from ∼ 10 mPa.s to ∼ 5 x 10^5^ mPa.s) ^26^. By packing micron-scale granules of hydrogels swollen in liquid LB (Luria-Bertani growth medium) beyond their jamming concentration, we manufacture a soft, semisolid 3D matrix with the key physical properties of mucus — especially the porosity (*ψ*), shear moduli (*G’* and *G’’*), and viscosity (*η*) (**Fig. 1d-1f, Extended data Fig. 2b, and 2c**). The nanometer-scale internal mesh sizes of the polymer chains forming each hydrogel granule allow unimpeded uniform diffusion of nutrients and small molecules throughout the bulk medium^27–29^. The microparticles themselves pack together to generate micron-scale inter-particle pore spaces (*ψ*), which provide a porous and internally disordered matrix for the bacteria. Within this, bacteria can move by exploring the tortuous pore spaces, as well as achieve 3D colony growth by pushing aside the microparticles to locally and reversibly deform the matrix. Importantly, our 3D microgel growth media retain soft solid-like bulk properties under low shear, while being reversibly fluidizable upon application of shear rates higher than the yield stress limit^30,31^—which is orders of magnitude lower than the internal turgor pressure of typical bacterial membranes. This allows bacterial colonies to internally rearrange the 3D matrix during growth without altering its material properties, allowing us to reproducibly maintain stable mechanical regimes over long-term culture. Using these platforms, we ask how two different degrees of confinement—representing both extremes of the physical regimes commonly observed for naturally occurring mucus—alter bacterial growth.

**Figure 2:**
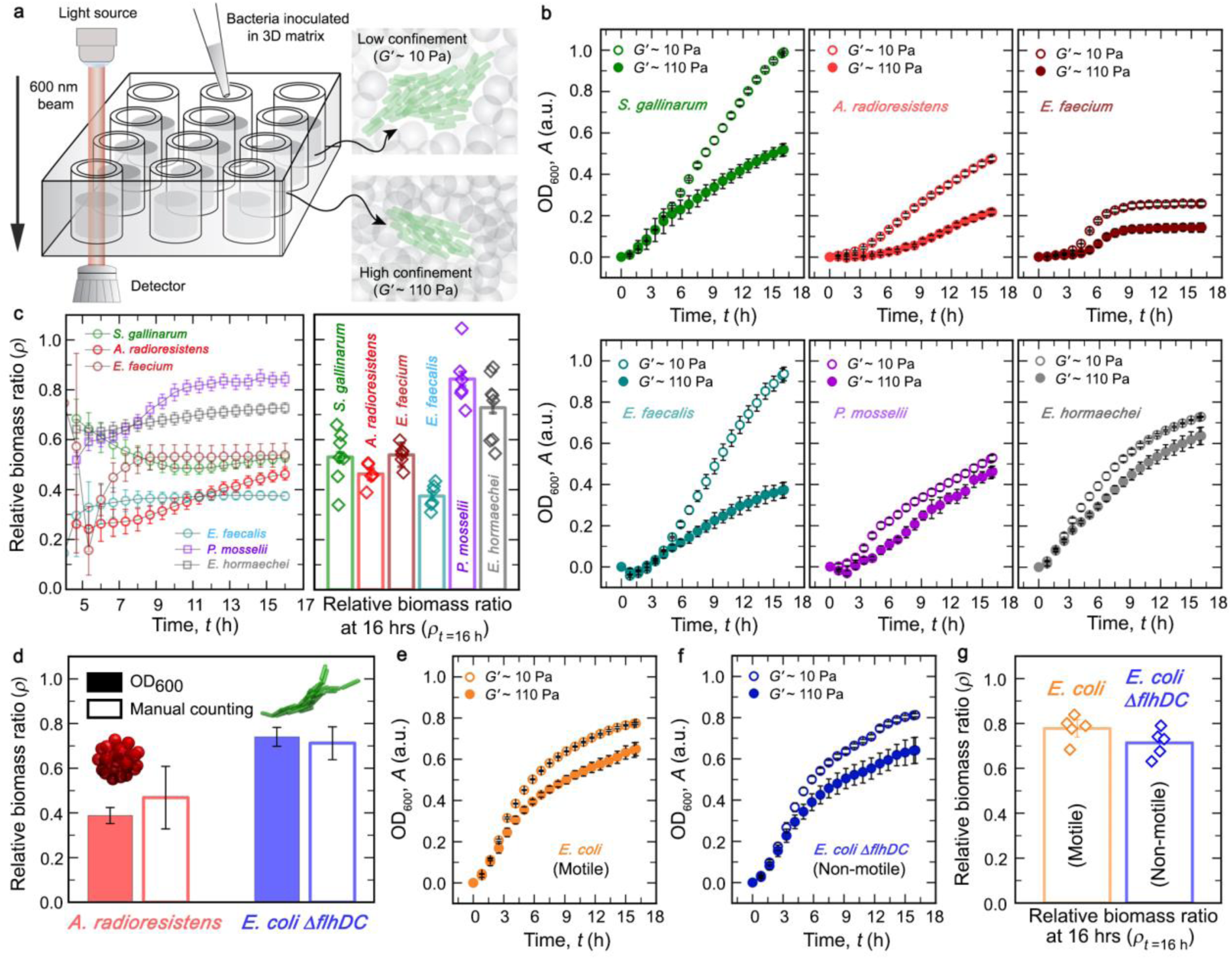
Change in confinement alters bacterial population growth. (a) We use absorbance-based optical density measurements as an indicator of increase in biomass over time to quantify bacterial growth in the 3D microgel media. (b) Representative absorbance-based growth curves over 16 hours indicating confinement-dependent growth dynamics for all bacterial strains (mean +/- s.d., n = 3 technical replicates); *G’* ∼10 Pa represents low confinement matrix and *G’* ∼ 110 Pa represents high confinement matrix. (c) Relative biomass ratio values for all bacterial strains across all time points, showing that higher confinement demarcates the different microbial species into two distinct sub- groups, indicated by bar graphs representing the plateau value of the relative biomass ratio at 16 hours (mean +/- propagated error, which is calculated as described in the statistical analysis, with biological replicates n = 7 for *S. gallinarum*, n = 5 for *A. radioresistens*, n = 6 for *E. faecium*, n = 6 for *E. faecalis*, n = 6 for *P. mosselli*, and n = 8 for *E. hormaechei*). (d) Absorbance measurements from 3D growth media mirror the actual microbial load, determined by measuring volumetric bacterial density via direct counting of cells using 3D confocal microscopy (also see Extended data Fig. 2e). Both of these measurements yield comparable relative biomass ratio after 16 hours of growth (mean +/- propagated error, which is calculated as described in the statistical analysis; for *A. radioresistens* n = 3 technical replicates for absorbance measurements; while for the manual counting 317 cells were pooled from 3 unique micrographs for the low confinement gel, and 198 cells were pooled from 4 unique micrographs for the high confinement gel – all sourced from the same biological replicate; and for the non-motile *E. coli*, n = 4 technical replicates for absorbance measurements; while for the manual counting 1163 cells and 828 cells were pooled from 5 unique micrographs each for the low confinement gel and high confinement gel, respectively – all sourced from the same biological replicate). (e-g) Motility does not confer significantly greater fitness benefits under confinement, as tested using a motile and non-motile strain of *E. coli* (the latter carries a deletion in the flagellar protein-encoding gene *flhDC*), which show no significant differences between the growth curves (mean +/- s.d., n = 3 technical replicates) or the relative biomass ratio (mean +/- propagated error, n = 5 biological replicates).

### 2.2 Bacterial growth varies with confinement

The 3D growth medium comprises a low solid fraction, which lends it high optical transparency allowing us to directly visualize bacterial growth under physical confinement. Leveraging this physical property, we homogeneously disperse bacteria in both low confinement and high confinement matrices and employ standard optical density-based absorbance measurements for assaying the temporal change in bacterial biomass (**Fig. 2a**). Optical density measurement (*A*) in these matrices is linearly proportional to the volumetric density of bacterial cells (*λ*) as obtained by direct counting from micrographs. Our volumetric counting method is quantitively robust as it correlates well with the conventional colony forming units (CFU)-based assay for identical samples (**Extended data Fig. 2e, 2f, and 2g**). Interestingly, we observe a decrease in the overall biomass production for all bacterial strains with an increase in confinement (**Fig. 2b and Extended data Fig. 3**). Prior reports suggest that the growth of bacteria is altered under elevated hydrostatic pressure—well above the turgor pressure—which may lead to deformation of the cell shape and induce stress response ^32,33^. However, since the elastic modulus of the microgel growth media is approximately three orders of magnitude lower than the internal turgor pressure of bacterial cells^34^, the observed reduction in overall biomass production is likely not due to compressive deformation of the cell body. To rule out aberrant effects of elevated osmotic stress on bacterial growth, we culture bacteria in liquid LB broth doped with varying amounts of the bioinert polymer PEG-1000 that induces an increase in osmotic pressure up to 100-fold higher than that exerted by our highest degree of 3D confinement. We find minimal effects on bacterial growth (**Extended data Fig. 4a)**. We further verify that our 3D matrices do not significantly affect cellular morphology. We negate the possibility of body deformation under 3D confinement by morphometrically analysing the cells; in both low and high confinement matrices, cell morphology remains identical, indicating that growth decrease under higher confinement is not due to decrease in cell sizes (**Extended data Fig. 5a, 5b, and 5c)**. Further, we show that the observed growth patterns cannot be attributed to cell death by quantifying the cell viability in the microgel growth medium. These results show that growth under high confinement does not detrimentally affect the proportion of viable cells in the population (**Extended data Fig. 5d and 5e).** To further ensure that an increase in confinement does not cause detrimental effects on cellular physiology, we also culture a genetically modified *Escherichia coli* strain— engineered to visually report both the genome’s physical organisation using HU (a DNA-binding protein) tagged to GFP, as well as induction of SulA-driven DNA damage responses — in 3D microgel growth media of both low and high degrees of confinement (**Extended data Fig. 5f**). We observe that increase in physical confinement neither alters the HU-GFP localization pattern (i.e., no observable change in the genome’s spatial arrangement ^35^) nor induces SulA expression (indicating DNA damage responses are not triggered ^36^). Hence, the decrease in biomass production under higher confinement is likely not due to altered growth patterns such as those triggered by DNA damage-induced stress responses.

**Figure 3:**
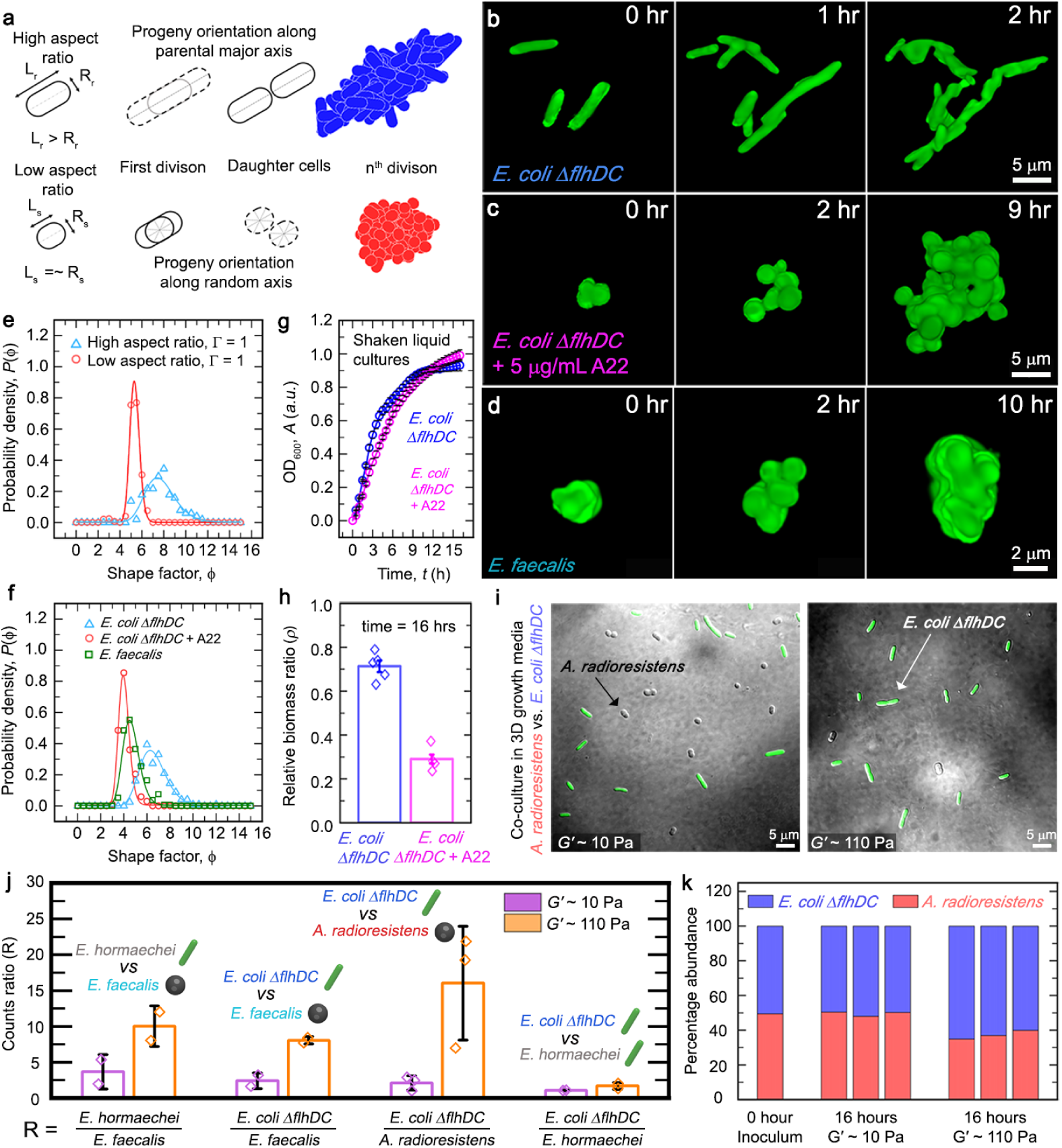
Bacterial morphology determines colony architecture and competitive outcomes. (a) We perform agent-based simulation of 3D colony growth for both high and low aspect ratio cells. In these simulations, progeny of high aspect ratio cells exhibits a directional bias along the parental axis, whereas those of low aspect ratio cells are more flexible to grow along any random axis. (b) Rod-shaped high aspect ratio *E. coli* cells give rise to elongated colonies under confinement as they proliferate by pushing their progeny outwards (similar phenomena have been observed for at least three biological replicates; representative micrograph has been shown from a replicate with n >= 40 distinct individual colonies imaged). (c) Treatment with 5 μg/mL of the MreB-perturbing compound A22 produces approximately spherical *E. coli* cells, which result in a rounded colony architecture (similar phenomena have been observed for at least three biological replicates; representative micrograph has been shown from a replicate with n >= 10 distinct individual colonies imaged). (d) Naturally spherical *E. faecalis* cells, stained here with Calcein-AM Green, show division and colony formation patterns similar to that predicted by the simulations for low aspect ratio cells as well as the drug-treated spherical *E. coli* cells (similar phenomena have been observed for at least three biological replicates; representative micrograph has been shown from a replicate with n >= 50 distinct individual colonies imaged). Micrographs in (b), (c), and (d) are generated by capturing the front view of a 3D volumetric projection. (e and f) Quantitative analysis of colony morphology using the shape factor (ratio between perimeter and square root of the area obtained from 2D projections of colonies) shows a significant difference between high and low aspect ratio cells, for both computational simulations (n ≥ 200 snapshots, from 10 simulations) and experiments (n ≥ 200 individual colonies). (g) Shaken liquid cultures in LB media for both rod-shaped *E. coli* and A22-treated spherical *E. coli* cells do not show any significant differences in their overall growth (mean +/- s.d., for n = 3 technical replicates). (h) Relative biomass ratios for rod-shaped and spherical *E. coli* indicate that rod-shaped cells are less affected than spherical ones under increased confinement, i.e., have ρ values closer to 1 (mean +/- propagated error, which is as described in the statistical analysis, with biological replicates n = 5 for rod-shaped non-motile E. coli and n = 4 for A22-treated spherical non-motile E. coli). (i and j) Direct visualization and endpoint counting of cells from 3D co-culture assays including both low aspect ratio (A. radioresistens or E. faecalis) and high aspect ratio cells (motile E. hormaechei or non-motile E. coli) show that the latter enjoy a growth advantage under increased confinement, at the expense of the former. However, competitive cultures between two high aspect ratio strains do not show significant enrichment of either strain (mean +/- s.d., for E. hormaechei vs E. faecalis, n = 2 biological replicates, each with >= 400 individual cells pooled from >= 10 unique micrographs for each gel type; for non-motile E. coli vs E. faecalis, n = 2 biological replicates, each with >= 300 individual cells pooled from 5 unique micrographs for each gel type; for non-motile E. coli vs A. radioresistens, n = 3 biological replicates, each with >= 180 individual cells pooled from 10 unique micrographs for each gel type; for non-motile E. coli vs E. hormaechei, n = 2 biological replicates, each with >= 130 individual cells pooled from 10 unique micrographs for each gel type). (k) Analysis of genomic DNA extracts from similar 3D co-culture setups confirm that growth of high aspect ratio bacteria are more robust to increase in physical confinement compared to that of low aspect ratio cells (n = 3 biological replicates).

**Figure 4:**
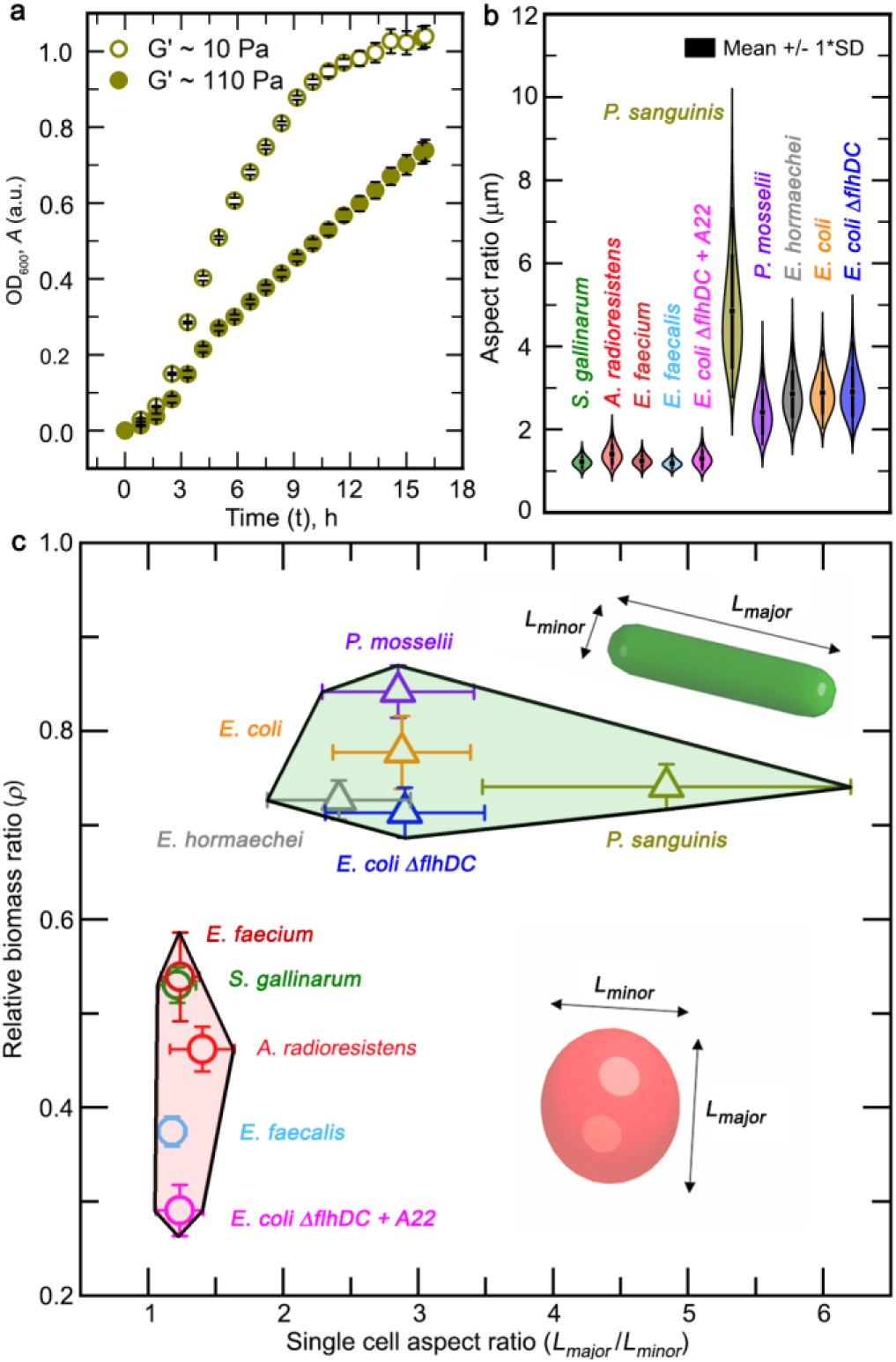
Single cell morphology enables physical confinement to selectively favour elongated colony architecture. (a) Growth assays indicate that *P. sanguinis* performs at par with rod-shaped bacteria across two different degrees of confinement (mean +/- s.d., n = 3 technical replicates. (b) Measurements of the aspect ratios of bacterial cell bodies for all ten strains used in this study, measured from samples in the late exponential phase (n ≥ 80 individual cells represented as lognormal distribution, with boxes indicating the means, and bounds of the boxes indicating +/- s.d.). (c) Microbial growth under confinement, as described using a phase space defined by the relative change in biomass production under increased confinement (the relative biomass ratio) vs. single cell morphology (mean +/- propagated error for the relative biomass ratio and mean +/- s.d. for the single cell aspect ratio).

To quantify the impact of confinement on growth, we normalize the biomass achieved at each timepoint in the high confinement matrix (*A*_high|16h_) to that in the low confinement matrix (*A*_low|16h_) for each strain, which gives the relative biomass ratio (*ρ*). This parameter initially varies with time, before achieving a plateau over twelve to sixteen hours for all strains, indicating the existence of a general, non-species-specific effect of confinement on bacterial growth (**Fig. 2c**). To focus on these effects at a stable time point, we quantify the endpoint biomass ratios at the sixteen-hour mark and observe that strains cluster into two well-separated groups—those which perform poorly under increased confinement, and those which remain relatively less perturbed (**Fig. 2c**). For additional rigor, we directly benchmark measurements of endpoint absorbance ratios against volumetric density of bacterial cells as obtained by microscopic counting from the same samples. Importantly, we perform these confirmatory tests for cells possessing two different shapes—approximately spherical and rod-like—thereby establishing the relative biomass ratio (*ρ*) as a generalised quantitative measure of change in bacterial biomass in 3D matrices (**Fig. 2d**).

A comparison of the various strain-specific morphological attributes listed in Fig. 1b reveals motility and cell body shape as distinguishing features of the two clusters observed in Fig. 2c. Since motility can conceivably allow better dispersion of bacteria in the 3D matrix ^24^ — thereby improving overall population-level growth through better nutrient accessibility and lesser local crowding—we first test the impact of motility on enhancing growth by employing both a motile (wild-type) and non-motile strain of *E. coli* (harbouring a deletion of a flagella-encoding gene). Surprisingly, both strains perform similarly under different degrees of confinement as captured by their relative biomass ratio (*ρ*) (**Fig. 2e-2g, Extended data Fig. 5g, and 5h**), invalidating motility as the dominant feature mediating the growth patterns observed under increased confinement. Consequently, we revisit the set of species-specific morphological attributes and note that while all poorly-performing strains comprise low aspect ratio (approximately spherical) cells, the better-performing strains have high aspect ratio cells (rod-like). These effects are not due to different biological species possessing intrinsically different growth rates. Although we observe that the absolute growth of certain coccoidal species is higher than some bacilli in the low confinement matrices, the relative biomass ratio metric specifically captures the performance of a given strain across two different degrees of confinement and is hence agnostic to how any other species fares. Using this, we observe a specific trend favouring a particular class of bacteria over others only when analysed along the shape axis. This observation forces a significant realignment of our perspective on growth under confinement, implicating cellular morphology as the attribute critical to growth success under spatial constraints.

### 2.3 Single-cell morphology dictates colony shape

Unlike growth on homogeneous 2D interfaces or in well-mixed liquid media, growth under physical confinement imposes spatial constraints and consequent nutrient limitations on bacterial colonies. To determine how different single-cell morphologies affect colony architecture, we employ an agent-based simulation using the CellModeller platform ^37^, maintaining all growth parameters constant and only altering the cellular aspect ratio. We simulate growth undertaken by low aspect ratio (approximately spherical) and high aspect ratio (rod-shaped) cells (**Fig. 3a and Supplementary video 1**). While most high aspect ratio bacilli (rod-shaped) cells such as *E. coli* exhibit cell division patterns that produces progeny cells oriented along the parental cell’s longitudinal axis, several low aspect ratio coccoid (spherical) species show planes of division that are either randomly or perpendicularly positioned to previous division planes, which allow a higher degree of freedom for the orientation of progeny cells following division ^38^. We incorporate these patterns in our simulations by implementing that high aspect ratio bacteria divide and produce progeny oriented along the length of the parent cell, whereas low aspect ratio bacteria produce progeny oriented along a randomly chosen direction. Under these conditions, striking differences emerge, wherein the high aspect ratio bacteria generate elongated colonies, while the low aspect ratio bacteria give rise to rounded 3D colonies. We further characterise these using a parameter termed the shape factor (*ϕ*), defined as the ratio between the perimeter and square root of area for the projection of a given colony. A low value for the shape factor (*ϕ*) indicates that the colony has a relatively lower exposed surface area as opposed to the enclosed volume. We find that the low aspect ratio bacteria predominantly organise into colonies with a low shape factor, and the high aspect ratio bacteria form elongated, high shape factor colonies (**Fig. 3e, Extended data Fig. 6d, and 6e**). Hence, single cell morphology appears to strongly dictate overall colony architecture.

We experimentally test this by live imaging the 3D colony growth of non-motile rod-shaped *E. coli* under high confinement matrices and observe patterns consistent with our computational model. Rod-shaped *E. coli* cells undertake volumetric growth along the parental axis, and push their progeny cells outward, leading to elongated colony architecture (**Fig. 3b, Extended data Fig. 6a, and Supplementary video 2**). Next, to test these patterns using a spherical bacterial species, we fluorescently label *E. faecalis* cells using Calcein-AM Green. Time lapse imaging shows that these coccoid cells also follow a colony growth mode similar to that predicted for low aspect ratio cells by the simulations, by organizing into well-rounded colonies (**Fig. 3d, Extended data Fig. 6c, and Fig. 8d)**. Further, we directly alter the morphology of non-motile *E. coli* using sub-MIC dosages of the MreB-perturbing compound A22 ^39^. Sub-lethal concentrations (5 μg/ml) of the drug induce aberrant division, rendering the rod-shaped *E. coli* into spherical cells. Again, as predicted by our computational model, the A22-treated low aspect ratio cells assume a rounded colony morphology (**Fig. 3c, Extended data Fig. 6b, and Supplementary video 2**). By quantitatively analysing the shape factor (*ϕ*) of the max projected fluorescent micrographs of colonies formed by rod-shaped and spherical cells, we find both the experimental and computational data to be in good agreement, with the single cell morphology being strongly implicated as the key determinant of colony-level organisation under 3D growth conditions (**Fig. 3f**).

### 2.4 Confinement selectively favours rod-shaped cells

A comparison of the relative biomass ratio (*ρ*) for rod-shaped and spherical *E. coli* reveals that alteration in shape severely compromises growth success under increased confinement, mirroring the trends exhibited by high and low aspect ratio gut-derived bacteria. Under similar concentrations of drug treatment (5 μg/ml A22), no significant growth differences are observed in liquid media. (**Fig. 3g, 3h, Extended data Fig. 6f, and 6g**). In fact, the relative biomass ratio of the spherical *E. coli* is quite similar to that of the naturally low aspect ratio cells (approximately *ρ* ≤ 0.5). Together, these observations highlight how cellular morphology enforces differences in the colony architecture of rod-shaped and spherical cells during growth under confinement. We further find that these trends depend only on the cellular shape rather than other environmental variables. To test this hypothesis, we first alter the nutrient composition for the 3D matrices from rich LB broth to minimal media (M9) supplemented with glucose and amino acids (**Extended data Fig. 7a and 7b**). When cultured in the minimal media matrices, high aspect ratio *E. coli* and low aspect ratio *A. radioresistens* mirror the relative biomass ratio trends observed in LB-based matrices, indicating that the relative biomass ratio patterns are independent of nutrient composition (**Extended data Fig. 7c-7e**). The above results lead to the prediction that all else being equal, rod-shaped bacteria should have a growth advantage over spherical bacteria under higher confinement. We additionally verify this using numerical calculations of bacterial colony growth, wherein we consider two different colony architectures—ellipsoidal and spherical— formed by high aspect ratio rod-shaped bacteria and low aspect ratio spherical bacteria, respectively. The effective local concentration of a nutrient inside a 3D bacterial colony is a result of two processes: replenishment of small nutrient molecules via unimpeded diffusion and depletion of nutrients via consumption. The competition between consumption and diffusion sets a depth of 10 µm from the surface of a 3D bacterial colony beyond which the nutrient is completely depleted ^20^. Factoring in this previously established limit on effective nutrient availability inside a 3D bacterial colony—which sets a limit on the actively proliferating fraction of a given 3D bacterial colony—we estimate the proportion of actively growing bacterial cells in colonies of either geometry with different sizes (**Extended data Fig. 6h and 6i**). Ellipsoidal colonies comprising of rod-shaped cells retain a higher proportion of cells in the actively growing zone as opposed to spherical colonies of comparable sizes. Together, the above results indicate that high aspect ratio bacteria produce elongated colonies, which allow better nutrient access in a 3D matrix, thereby enabling superior growth under increased confinement.

To test whether this effect provides a competitive advantage to high aspect ratio cells in a mixed population, we carry out co-culture assays that combine both high aspect ratio and low aspect ratio cells across two different degrees of confinement (**Fig. 3i-3k, Extended data Fig. 8a, 8b, and 8c**). For this, we prepare an inoculum with equal parts of two different bacterial species that we homogeneously disperse in both low and high confinement 3D growth matrices. More specifically, we perform competitive co-culture assays between the following five combinations: (i) high aspect ratio motile *E. hormaechei* and low aspect ratio non-motile *E. faecalis*, (ii) high aspect ratio non-motile *E. coli* and low aspect ratio non-motile *E. faecalis*, (iii) high aspect ratio non-motile *E. coli* and low aspect ratio non-motile *A. radioresistens*, (iv) high aspect ratio motile *E. hormaechei* and high aspect ratio non-motile *E. coli*, and (v) high aspect ratio motile *P. mosselii* and high aspect ratio non-motile *E. coli*. Following sixteen hours of growth, we mechanically disrupt the colonies in each matrix into homogeneously dispersed single cell suspensions. Subsequently, we tally the number density of each type of bacteria using microscopy to obtain the relative strain ratios for each competing pair. As expected, the relative proportion of high aspect ratio cells in the population significantly increases with an increase in confinement, at the cost of a large decrease in the abundance of low aspect ratio cells (**Fig. 3i, 3j, and Extended data Fig. 8a**). Interestingly, an increase in confinement does not alter the relative abundance of two high aspect ratio species in direct competition (**Fig. 3j and Extended data Fig. 8b**). Parallelly, we also perform bulk genomic DNA extraction and quantify the relative abundance of each strain using 16s rRNA amplicon sequencing. Our bulk genomic analysis qualitatively mirrors the trends observed with microscopic tallying of the number density: despite identical nutrient media, rod-shaped bacteria gain a substantial advantage over spherical bacteria under high confinement (**Fig. 3k and Extended data Fig. 8c**). These results thus validate high aspect ratio morphology as a beneficial attribute under spatially confined growth. Of course, in many ecological settings, inter-species interactions are also heavily influenced by cross-feeding, cooperative and antagonistic behaviours, fluctuations in the environment, resistance to invasion, predation, and chemical cross-signalling ^1–6^. Our platform presents the capability to conserve the heterogeneous chemical and behavioural landscapes introduced by such variables, and study how physical confinement acts on top of these processes. This is because the porous nature of our matrix allows for unimpeded small molecule diffusion^27–29^ regardless of the degree of confinement and thus ensuring that modulation of the mechanical regime can occur independent of all other factors.

In nature, cells exhibit morphological alterations under specific environmental conditions. For instance, uropathogenic *E. coli* transition between coccoid and rod-like morphologies during infection progression ^40^. These transitions are also subject to chemical signalling and nutrient availability. These are best illustrated by examples such as those of *V. cholerae*, which shows a curved rod-to-sphere transition in response to both low temperatures or L-arabinose treatment, as well as *L. monocytogenes* that undergoes a similar transition during biofilm formation on surfaces ^41–44^. Such shape changes likely confer survival advantages under suboptimal conditions such as extreme environmental changes or nutrient starvation. However, it is also conceivable that a sphere-to-rod transition might prove particularly beneficial purely in terms of nutrient availability in porous 3D environments such as tissues and soil by promoting more efficient growth and proliferation. We test this hypothesis using *P. sanguinis*, a species wherein the spherical form can grow in an elongated fashion as a rod — similar to previous reports on *P. denitrificans* ^45,46^ (**Fig. 4a, Extended data Fig. 8e, 8f, and Supplementary video 3**). We assay the growth of *P. sanguinis* across two different degrees of confinement and find that it mimics the relative biomass ratio trend observed for high aspect ratio bacteria. This further validates that the elongated colony structure of high aspect ratio strains confers a relative advantage under increased degrees of confinement.

Thus, we conclude that the aspect ratio of individual cells is a key determinant of the relative biomass ratio under 3D confined growth. To test this, we measure the cell body morphology of all the different bacterial strains used in our experiments and plot their individual aspect ratio by measuring the major axis and minor axis of the cell body. Distribution of these aspect ratios captures two different populations, separated by their morphology (**Fig. 4b**). Superimposing this with our previous results on the relative growth ratios across two different degrees of confinement, we observe a remarkable compartmentalization of the different bacterial strains into two non-overlapping domains. This phase space—defined by single-cell morphology and consequent growth performance under elevated confinement— underscores the idea that with an increase in confinement, single-cell morphology is the primary determinant of growth success in complex environments (**Fig. 4c**). Effectively, we demonstrate that the selective pressure exerted by the physical microenvironment in terms of confinement-induced spatial constraints strongly favours high aspect ratio cellular morphology due to its efficient colony organisation.

## 3. Discussion

Our work provides experimental evidence implying that physical confinement can play a selective role in deciding bacterial growth fitness within their natural niches. High aspect ratio bacteria leverage growth anisotropy to form elongated colonies with higher surface area, allowing better nutrient accessibility as opposed to the spherical colonies formed by low aspect ratio bacteria. Increased confinement strongly selects high aspect ratio cells for more efficient growth — a trend that holds even for high and low aspect ratio forms of the same bacterial strain, indicating a remarkable interconvertible behaviour governed solely by cellular shape. Importantly, we demonstrate the potential to use such constraints to alter bacterial community composition under biomimetic mechanical regimes. Thus, our results represent an important conceptual advancement towards experimentally modelling bacterial growth within complex natural habitats.

In microbial ecology, the present mechanistic understanding of predator-prey interactions, niche partitioning, and community organization largely relies on a framework comprising genetic mutations, behavioural patterns, and biochemical signalling ^47–51^. Notably, our platform does not preclude such chemical interactions. Instead, the porous and permeable nature of our 3D matrix presents an opportunity to capture the dynamics of a chemical signalling landscape coupled with different mechanical regimes, enabling a more comprehensive experimental recapitulation of natural niches. While the permeable nature of our 3D matrix allows unimpeded small molecule diffusion, several biological niches such as soil and aquifers resemble semi-permeable porous media with non-deformable, rigid physical barriers. We expect that the reduced nutrient availability and severe spatial restrictions in such contexts significantly restrict colony growth and organization. Recreating such regimes *in vitro* will require the development of semi-permeable 3D scaffolds. Further, within the temporal window of our assays, we do not observe significant contributions from either aberrant phenotypes or perturbed physiologies. However, environmental pressures in the form of nutrient deprivation, high mechanical stiffness, and antagonistic interspecies interactions are known to induce the expression of stress genes, alter metabolism, trigger extreme phenotypes (such as filamentous growth), and favour increased mutational rate ^52–57^. Such factors are likely to affect long-term bacterial growth— hence, future work that investigates bacteria under similar conditions over extended durations could provide interesting insights into the long-term effects of confinement on bacterial communities. Presumably, our system can be engineered to recreate such effects by altering the nutrient media composition and tuning the mechanical stiffness of the matrix. We speculate that such environmental constraints will selectively favour a subset of adaptive mutations with the potential to alter evolutionary trajectories.

Furthermore, it has been reported that spatial patterning within mixed microbial communities can be strongly driven by single cell morphology ^58^. This suggests an interesting direction for future exploration. Given the selective pressure enforced by elevated physical confinement favouring rod-shaped bacteria over spherical ones, how do existing spatial patterns get altered when the microenvironmental mechanics change? Conversely, does physical confinement alter the evolution of spatial architecture within mixed microbial communities such as biofilms? Finally, our work highlights the need to explore the role of physical confinement in diverse niches that may select for varied life history strategies. For instance, short-term rapid growth and expansion are not universally preferred traits. In adverse conditions such as in the presence of toxins or predators when long-term cell survival is prioritized, a transient shift towards quiescence or dense collective organisations with minimally exposed surface areas may be more advantageous ^59–63^. Such conditions and strategies may override the selection on cell shape imposed by physical confinement and should be explored further.

Current descriptions of microbial growth invoke either genetic mutations or biochemical signalling as the primary mechanisms via which bacteria respond to their microenvironment. By contrast, we argue that the spatial constraints imposed by their microenvironment exert a more general effect, irrespective of specific organismal biology. Our conceptual framework can be employed to generate coarse-grained predictions of population dynamics under different biophysical regimes. Understanding how bacteria grow in complex, disordered 3D environments such as tissues, mucus, and soil, is also central for combating antibiotic resistance, improving agricultural practices, and bioremediation — underscoring the high-value practical applications of our work.

## 4. Methods

### 4.1 Preparation and Characterization of 3D growth media

We prepare the 3D matrices by dispersing dry hydrogel granules of Carbopol C-980 (Lubrizol) at either a 0.50% or 0.85% (w/v%) concentration in 2% (w/v %) liquid LB (Sigma, LB Lennox) media. While preparing minimal media matrices, we replace LB media with M9 media (BD Difco, M9 Minimal Salts) supplemented with 0.2% glucose and 0.2% milk protein hydrolysate as a source of amino acids. The hydrogel suspension is vigorously stirred for at least 2 hours to ensure complete hydration of the dry granules. Since the Carbopol C-980 is a negatively charged hydrogel, we neutralize the pH of this suspension to 7.4 using 10N NaOH. At pH 7.4 the hydrogel granules or microgels swell maximally to an extent when individual granules jam against each other forming a transparent and disordered solid matrix. We determine the viscoelastic properties of jammed microgels using oscillatory shear rheology on an Anton Parr (MCR302e) rheometer with a roughened cone-plate measuring tool. We adjust the solid fraction of 3D growth media to achieve an elastic storage modulus range between 10 Pa to 110 Pa, that spans the measured range of biological mucosal samples. To measure the yield stresses and viscosity of the hydrogels, we apply unidirectional shear at varying rates while recording the shear stress response. As shown in Extended data Fig. 2b, our microgels behave as yield stress materials, which are reversibly fluidizable upon application of the threshold shear rate but retain a soft solid-like nature below this regime. Porosity of the microgels are determined using 200 nm fluorescent tracer particles. We track the thermal diffusion of these beads through the inter-particle pore spaces. Since the internal mesh sizes of the hydrogel granules are smaller than 100 nm, the beads are occluded from these regions. Hence, the beads only explore volumes that are also accessible to bacteria (which are up to 10-fold larger in dimension than the bead size). We disperse a dilute solution of the tracer particles in both low and high confinement matrices. Subsequently, we acquire images within a time interval of 17 ms using the 40X objective of an inverted laser-scanning confocal microscope (Nikon A1R HD25) to track their thermal diffusion-guided trajectory. Individual beads are tracked with sub-pixel precision using a custom-written MATLAB script based on the Crocker-Grier algorithm ^64^, and the mean-square displacement (MSD) is quantified as a function of lag time. Over short time scales, the particles freely explore the pore spaces. However, over longer time scales, their motion becomes constricted by the porous medium and its dead-end channels. This implies that the particle MSDs will increase linearly with time for unimpeded motions, while over longer time scales the pore boundaries will make the MSDs plateau. The characteristic smallest pore space dimensions explored by the particles is then given by the square root of these plateau values added to the particle diameter. We calculate the 1-CDF (complementary cumulative distribution function) of the measured pore spaces, which represents the overall pore size distribution within our 3D microgel growth media. Herein, the value of the 1-CDF at any given pore size represents the fraction of the pores which are larger than the given pore size. For instance, a 1-CDF value of 0.5 represents the 50th percentile of pore sizes. Similarly, a 1-CDF value of 0.05 denotes that 5% of the pores are larger than the corresponding pore dimension. (**Extended data Fig. 2c**). We also show that under the culturing conditions followed in this work, our microgel samples do not undergo any significant alterations to the rheological properties or bulk volumes. For this, we take the low confinement matrix, and maintain it at 30⁰C for up to 24 hours under humid conditions (**Extended data Fig. 2a).** This is similar to the assay set up for all our biological experiments. Further, we recover and pool together the microgel samples from these plates and carry out rheological measurement as well as direct weighing to establish that both the rheological properties and the overall sample volume remains conserved (**Extended data Fig. 2h, 2i, and 2j)**.

### 4.2 Isolation, *in vitro* culture, and characterization of bacterial strains

We isolate seven different microbial species from red flour beetles (*Tribolium castaneum*; the red flour beetles were collected by DA from a flour warehouse in India and have been maintained by DA for ∼ 10 years at NCBS) and identify these by Sanger sequencing of the 16S rRNA gene using the universal 16S primers 27F (5’-AGAGTTTGATCCTGGCTCAG-3’), and 1492R (5’-GGTTACCTTGTTACGACTT-3’). Following the alignment of both forward and reverse sequences, a homology search using NCBI BLAST is carried out to identify the closest bacterial species for each such amplicon (16S sequences provided for each isolate in **Extended data Table 1**). Primary isolates from these are cultured in 2% LB broth and cryopreserved as glycerol stocks (20% v/v). For each experiment, a shaken culture of 1% stock dilution in LB broth is grown at 30⁰C for ∼20 hours to achieve a stationary phase culture. From this, a 1% inoculum is added to fresh 3D growth media and homogeneously dispersed by gentle pipette mixing. To promote better growth in the case of *E. faecalis* and *E. faecium*, overnight cultures are directly set using 3D hydrogel media. We also employ two strains of *E. coli* – one, the wild-type motile strain, and the second carrying a deletion in the flagellar gene flhDC, that renders it non-motile. These *E. coli* strains were a kind gift from Prof. Robert Austin, of Princeton University, which constitutively express cytoplasmic green fluorescent protein (GFP) and have been previously used to study bacterial motility in 3D porous media^24,27^. For antibiotic treatment of *E. coli* with A22, we maintain a sub-MIC concentration of 1 μg/ml in LB and 5 μg/ml in 3D media. To characterize the bacterial morphology, we dilute samples using either LB or 0.5% 3D growth media and spread these on a flat agar pad (1% (w/v) agar in water) placed inverted onto a glass bottom dish. Images are acquired using a point-scanning laser confocal microscope. To measure the cellular morphology, we segment micrographs in ImageJ and use in-built functions to quantify the cell body area, length, and aspect ratio.

We ascertain the suitability of LB broth as a nutrient medium for all bacterial strains by assaying their growth in shaken liquid cultures (**Extended data Fig. 1**). We separately assess the growth performance of these strains under non-shaken culturing conditions in liquid LB, results from which show that static culturing conditions in liquid media lead to less efficient growth for most microbial strains used in our study (**Extended data Fig. 4b)**. While shaken culture ensures well-mixed condition for uniform nutrient and oxygen availability throughout the bulk culture volume, in non-shaken culture progressive densification may occur over time due to cells settling under gravity. In contrast, physical confinement manages to maintain an initially homogeneous distribution of bacterial density throughout the bulk 3D volume of the material, while nutrient availability is also similar throughout due to the freely permeable and porous nature of the microgel matrix. Hence, both shaken and non-shaken culturing conditions are fundamentally different from the conditions encountered by bacteria embedded in our 3D matrix and hence growth performance in these conditions should not be compared.

We also quantify the viability of cells grown in the high confinement matrix. In all samples, we use propidium iodide (fluorescence in red) as a marker for dead cells, since this compound stains cells with compromised membrane integrity. We also use Calcein-AM Green as a marker for viable cells in the case of *E. faecalis* which shows good uptake of this compound. For the *E. coli* strains, we use the constitutively-expressed green fluorescent protein signal as a marker for viable cells. For viability testing, we culture different bacterial strains in the high confinement matrix for 24 hours, following which we dilute the samples in liquid LB, vigorously mix using vortexing to break apart colonies into single cells under mechanical shear, and incubate with the appropriate staining solutions, before spreading them on flat agar pads for image acquisition. To test for genomic DNA damage-triggered responses, we use *E. coli* strains carrying a constitutively-expressing GFP-tagged histone protein (HU-GFP) that marks the DNA, along with an mCherry-tagged DNA damage response-triggered protein (SulA-mCherry), received as a kind gift from Dr. Anjana Badrinarayan of NCBS. These are cultured for 24 hours in microgels of different stiffnesses and imaged using a confocal microscope. As an additional control, we induce DNA damage in an exponential-phase culture of bacterial cells cultured in liquid LB by treating with 5 μg/ml of the DNA damage-inducing drug MMC, which exhibits strong SulA-mCherry expression.

### 4.3 Growth assays

We assay microbial growth in 3D growth media using absorbance-based optical density measurements at 600 nm. Samples prepared as described above are dispensed as 200 μl replicates into 96-well plates, and readings are acquired at 10-minute intervals for 16 hours using a multi-mode plate reader (Varioskan Lux) with temperature control (set to 30⁰C). All data are normalized by subtracting the initial timepoint value (except for antibiotic-treated samples, where we normalize using the lowest absorbance value to account for initial cell death). This gives a measure of the net increase in biomass over time. For 3D co-culture assays, all parameters are maintained identical to the above, except the initial inoculum, wherein both competing strains are added in similar proportions by normalizing the optical density of overnight-grown cultures. To maintain identical culturing conditions as followed for the individual growth assays, we dispense these samples in a 96-well plate and grow them for 16 hours at 30⁰C. Following this, we pool together all the technical replicates for each gel type and homogeneously mix the sample. From this, a 20 μl aliquot is used for direct bacterial counting, with the rest being processed for genomic DNA extraction (described below). For growth experiments where we vary the increase in osmotic pressure, we use varying amounts of the inert polymer PEG1000 doped in liquid LB. Here, we maintain the elevation in osmotic pressures to levels similar to the high confinement matrix (∼110 Pa), as well as 10-fold and 100-fold higher. The osmotic pressures of these solutions are approximated using the equation *<u>icRT</u>*, where *π* represents the osmotic pressure, *i* denotes the van’t Hoff index, *c* is the molar concentration of solute, *R* is the ideal gas constant, and *T* is the temperature.

### 4.4. Direct microscopic counting of single cells

Briefly, we set up growth curve assays for the non-motile *E. coli ΔflhDC*, as described above, in multiple triplicate sets, for both low confinement and high confinement matrices. At specific time points separated by three-hour intervals, we obtain an optical density measurement for each set. Each individual replicate belonging to this set is harvested, mechanically sheared to break up colony-like clumps into single cells and diluted using LB (**Extended data Fig. 2d**). We scale the dilution factor for each timepoint following the first one to achieve approximately the same final density of bacterial cells. These samples are then mixed with microgel (to prevent cells from settling during the counting procedure) in a 1:1 ratio, following which 200 μl of this sample is dispensed into a glass-bottomed 35 mm dish and fixed 3D volumes from each are imaged using a confocal microscope. The total number of bacteria is tallied using automated thresholding and segmentation functions available in ImageJ, as well as the 3D Cell Counter plugin. Using this, we calculate the number of bacteria per mL from the original sample and plot this against the measured absorbance value. We observe a roughly linear relation between the microscopically counted number of cells and the optical density of the sample, confirming that absorbance values provide an accurate estimate of the total bacterial biomass in 3D growth media. We also directly benchmark the volumetric counting method against the conventional colony forming units (CFU)-based quantification method. Herein, we prepare samples of bacteria inoculated in the microgel growth media. These are diluted in liquid LB and imaged using confocal microscopy to obtain the volumetric cell density as described above. Following this, the same samples are serially diluted using liquid LB and plated on LB agar. Colonies resulting from these are counted the next day and tallied against the statistics obtained from direct volumetric counting. We find a good agreement between both these metrics, which together with the optical density vs. volumetric counting approach, establish that direct counting using microscopy is a quantitatively rigorous method which compares well against both optical density and CFU-based quantification methods.

To verify this for the relative biomass ratio metric, we set up growth curve assays for both *A. radioresistens* and the non-motile *E. coli ΔflhDC*, as described above, in both low and high confinement matrices. Immediately after the growth curve assay concludes, we pool together all replicate samples of each bacterial strain and each specific confinement type in separate microcentrifuge tubes. This is well-mixed to ensure homogeneous dispersion of the bacteria. 20 μl of this is added to 1 ml of 0.50% microgel (to prevent cells from settling during the counting procedure) and mixed thoroughly to break up the colonies into single cells. 200 μl of this sample is dispensed into a glass-bottomed 35 mm dish and imaged using a confocal microscope. From the images, individual bacteria are microscopically identified and tallied. We then compare the ratio of endpoint absorbances from 0.85% microgel to 0.50% microgel for both bacterial strains with the ratio of numbers obtained by direct counting for identical samples.

For the co-culture experiments, we prepare an inoculum with equal parts of both high aspect ratio cells (motile *E. hormaechei* or non-motile *E. coli*) and low aspect ratio cells (either *A. radioresistens* or *E. faecalis*), that we homogeneously disperse in both low and high confinement 3D growth matrices. Following sixteen hours of growth, we mechanically disrupt the colonies in each matrix into homogeneously dispersed single cell suspensions. Subsequently, we spread these on flat agar pads (in the case of the *E. coli* vs *A. radioresistens* combination, we also include one replicate with volumetric 3D cell counting) and tally the number density of each type of bacteria using microscopy to obtain the relative number of cells belonging to each strain type per sample.

### 4.5. Colony morphology experiments

For live-cell imaging of bacterial colonies, we use 35 mm plastic dishes with a central circular cavity (15 mm in diameter) sealed off with a glass coverslip for optimal quality imaging. Samples for assessing the colony morphology are prepared exactly as described above for the growth curve experiments. While the *E. coli strain* used in this study shows constitutive expression of the green fluorescent protein, we use the cell viability dye Calcein-AM Green to stain *E. faecalis* cells (for live imaging, the dye is directly added to the microgel growth media). 200 μl of well-mixed sample is dispensed into the glass-bottomed cavity of these dishes. Using a sterile coverslip, we scrape the top of this sample to ensure a smooth, flat layer. To prevent evaporative losses, we gently layer the top of the sample with 1.5 ml of mineral oil (Sigma). This oil is permeable to gases; hence the microgel-suspended bacteria are not cut off from oxygen supply. Plates prepared in this manner are placed in an incubator maintained at 30⁰C for 16 hours. Following this, we acquire z-stack images of the bacterial colonies using confocal microscopy. Colony morphology is analysed by segmenting 2D projections of the individual bacterial colonies, followed by calculation of the colony area and perimeter using in-built functions in Fiji-ImageJ. Using this, we calculate a parameter termed the shape factor, defined herein as the ratio between the perimeter and the square root of the area (for a circle, the shape factor equals 3.54). This quantity gives an estimate of how much colony shapes differ across various species and treatment conditions.

### 4.6. DNA Extraction and 16S rRNA amplicon sequencing

For genetic analysis of 3D media-grown and liquid (t_0_, initial inoculum) co-culture samples, we carry out genomic DNA extraction. 300 μl of each sample is incubated at 65°C for 2 hrs with 200 μl of Promega nuclei lysis buffer, followed by the addition of 200 μl Promega protein precipitation solution and 100 μl of chloroform to promote phase separation. The DNA-containing fraction is separated out by centrifugation, to which 300 μl isopropyl alcohol is added to precipitate the nucleic acids fraction. The resultant pellet is washed twice with 80% ethanol, following which we elute the genomic DNA in 40µl nuclease-free water. To determine the change in the relative proportion of the two co-cultured strains, we amplify the V3-V4 hypervariable region of the bacterial 16S rRNA gene using the modified Illumina 10N primer ^65^ and sequence amplicons on the Illumina Miseq platform (300x2 paired-end, obtaining ∼75000-140000 reads per sample). We analyze sequence data using the DADA2 pipeline ^66^ in R, which gives us the relative abundance of each strain in each sample.

We also note that the trends observed using 16S rRNA amplicon sequencing-based methods are largely qualitative in nature. This is primarily due to experimental limitations with designing efficient cell lysis protocols for both competing strains, as well as difficulties with optimal DNA extraction from cells embedded in 3D microgel matrices (wherein, cross-reactivity and charge interactions of the extraction reagents with the scaffold material cannot be entirely discounted). In this regard, the flexibility afforded by the matrix in terms of cell extraction and visualisation allow for parallel strategies such as direct counting of individual cell types, enabling a precise estimation of relative bacterial loads even from co-culture setups. However, direct visualisation and counting also comes with certain limitations. For instance, this strategy is only amenable for bacteria whose colonies can be sufficiently disrupted using mechanical shear without lysing the cells. Second, unless the competing strains are unambiguously distinguishable based on morphology (rod vs sphere) or visual contrast (e.g., fluorescently tagging/dye-based labelling or differential colorimetric staining), counting is not a reliable method. This limits its applicability for a select few combinations of bacterial strains. We believe that further work towards optimizing both 16S rRNA amplicon sequencing and visualisation-based approaches as well as developing hybrid workflows on a case-by-case basis will significantly improve the applicability of our 3D microgel matrix towards enabling a rigorous quantitative estimation of species-specific bacterial loads from mixed microbial communities cultured in vitro in 3D.

### 4.7. Simulating bacterial growth

We simulate 3D colony growth using a previously published agent-based platform (CellModeller^37^) which models bacterial cells as spherocylinders. We modify the original code by altering the aspect ratio of these objects by explicitly defining the radius and overall volume, which together adjust the cell’s length. Using this, we simulate 3D growth of either high-aspect ratio “rods”, or low-aspect ratio “spheres”. We trigger the cells to divide once they achieve a target volume greater than twice the defined volume for a single cell, with both resultant daughter cells starting off with the initial volume defined for a single cell. Further, we constrain the progeny cells produced by rods to maintain orientation along the parental axis, whereas the progeny cells produced by spherical cells are allowed to reorient in random directions. We also simulate different degrees of physical confinement by varying the environmental drag (Γ) on the growing bacterial cells.

### 4.8. Statistical analysis

For each strain of bacteria, we record the OD_600_ for a minimum of 4 sets of biological replicates in both low confinement and high confinements matrices. Each set of biological replicates for each type of matrix contains OD_600_ data from 2 to 4 technical replicates. For simplicity, consider the data from high confinement to have a mean *A*_*high*_ and standard deviation *dA*_*high*_, while for low confinement this would be *A*_*low*_ and *dA*_*low*_, respectively. This can be considered as two random variables *A*_*high*_ and *A*_*low*_, with errors of *dA*_*high*_ and *dA*_*low*_, respectively. Now, the relative biomass ratio (*ρ*) can be written as:

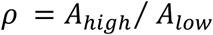

where *dρ* is the propagated error from the technical replicates of both *A*_*high*_ and *A*_*low*_. This can be formulated as:

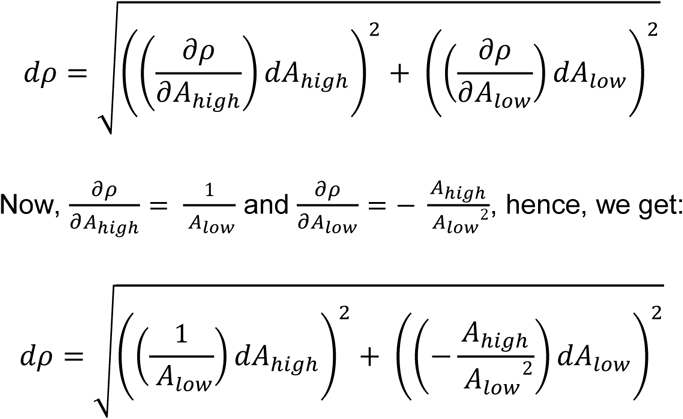

Multiplying and dividing R.H.S. by *ρ*, we get:

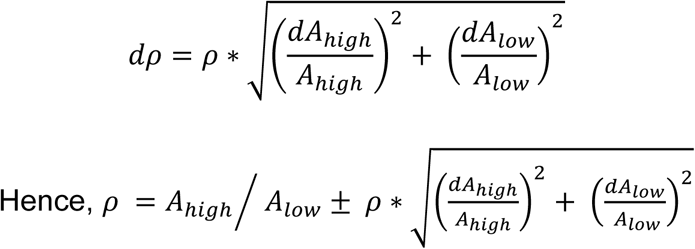

Now, consider the relative biomass ratios of any *n* such biological replicates, i.e., some *ρ*_1_, *ρ*_2_, … *ρ*_3_, from *n* independent experiments. The average of these will be given by:

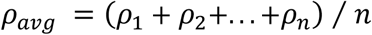

where, *dρ*_*avg*_ should represent the propagated error from all the individual relative biomass ratio values. This can be represented as:

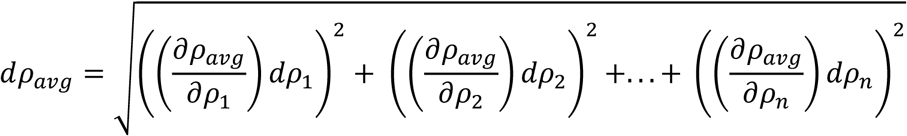

Herein, we know that 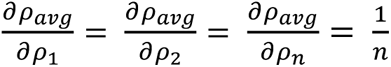, which reduces the above to:

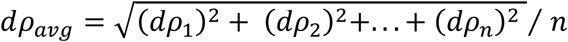

Hence, the net relative biomass ratio for *n* biological replicates, each with its own technical replicates, will be given by:

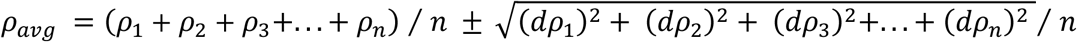

## Acknowledgments

T.B. and D.A. acknowledge intramural research grants from NCBS (Department of Atomic Energy Project 695 Identification No. RTI 4006). D.A. acknowledges a grant from the DBT/Wellcome Trust India Alliance (IA/I/17/1/503091) that also supported P.S. M.S. acknowledges personal support through the NCBS GS program. M.S. also acknowledges help from Saheli Dey, Kiruthika Kumar, and Manukrishna Manikandan for training on handling bacterial cultures and preparation of 3D growth media. We are grateful to Prof. Bob Austin for sharing the fluorescent *E. coli* strains and Dr. Anjana Badrinarayan for sharing the DNA damage-reporting *E. coli* strains. It is a pleasure to acknowledge the insights and helpful comments from Prof. Zemer Gitai and Prof. Sujit Datta, which help mould key concepts in this work. We also acknowledge the valuable inputs from Dr. Tim Rudge on computational simulations of bacterial growth, as well as helpful comments and discussions on the manuscript from Joanrose John Kattikaren. We thank Dr. Sneha Garge, Krithjgnan Bhardwaj, and Dhanush A for isolating bacterial strains from the flour beetle gut. We are grateful to Dr. H. Krishnamurthy and the Central Imaging and Flow Cytometry Facility, and the NGS facility at NCBS. We also acknowledge the NCBS common equipment facility for arranging the Varioskan multimode plate reader and the Anton Parr MCR 302e rheometer.

## Author contributions

T.B. conceptualized and supervised the study. T.B., M.S., D.A., and P.S. contributed towards the study design. M.S. performed the experiments, simulations, and data analysis, with help from M.G. P.S. characterized the gut-derived microbial strains and performed the genetic analysis. C.K. helped with the computational simulations setup. M.S. and T.B. wrote the manuscript with help from D.A. and C.K. T.B. acquired primary funding support. All authors provided critical feedback on the manuscript.

## Data availability statement

The 16S rRNA sequencing data of all beetle-derived primary isolates have been deposited in GenBank, accession codes for each of which are listed in Extended data Table 1. The co-culture data used in this study are available in NCBI SRA Bioproject accession no. PRJNA1169755.

Correspondence and requests for materials should be addressed to Tapomoy Bhattacharjee.

## Extended data

### Extended data figures

**Extended data Figure 1:**
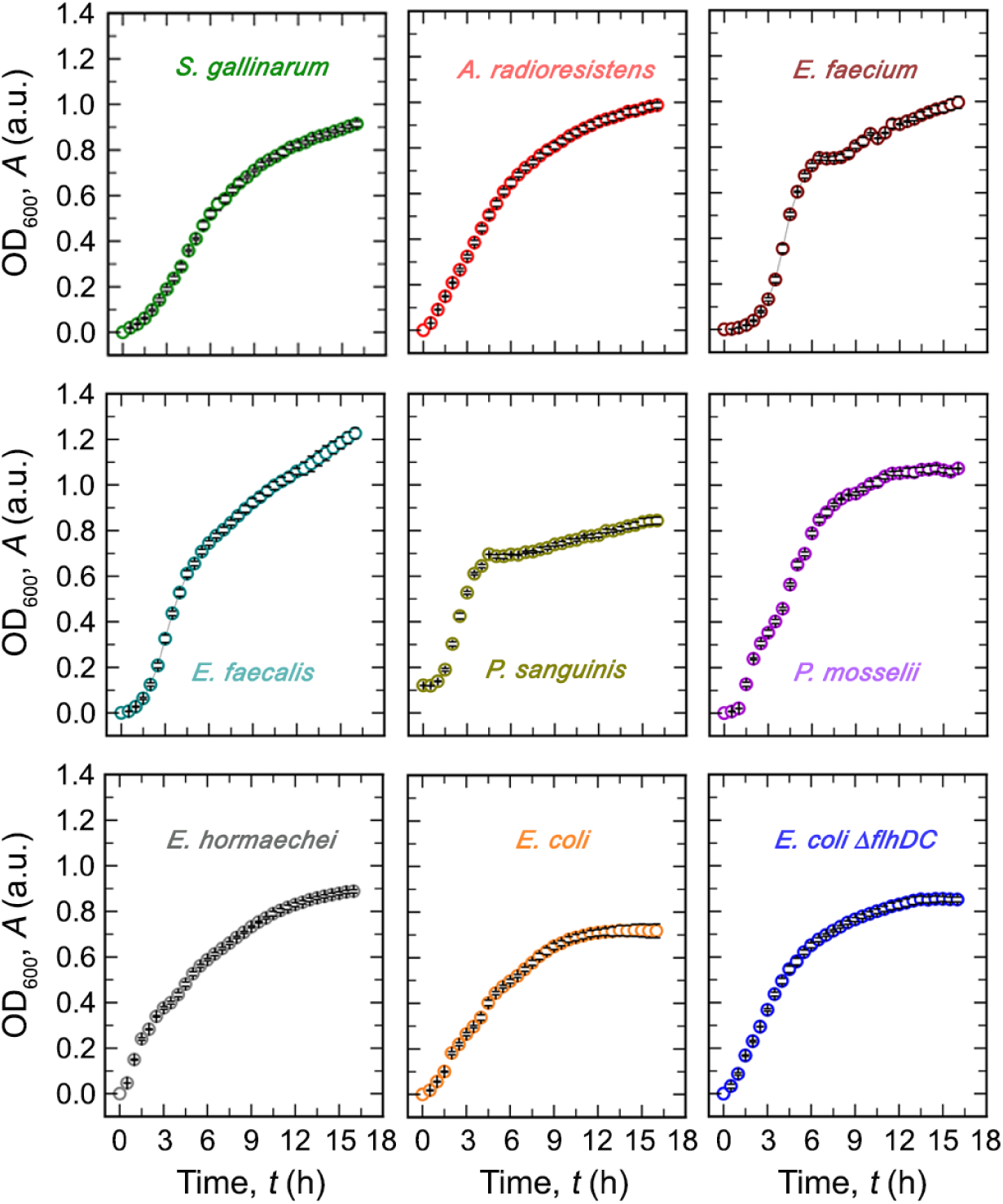
Absorbance-based growth curves for all bacterial strains in LB broth under continuously shaken culturing conditions (mean +/- s.d., for n = 3 technical replicates).

**Extended data Figure 2:**
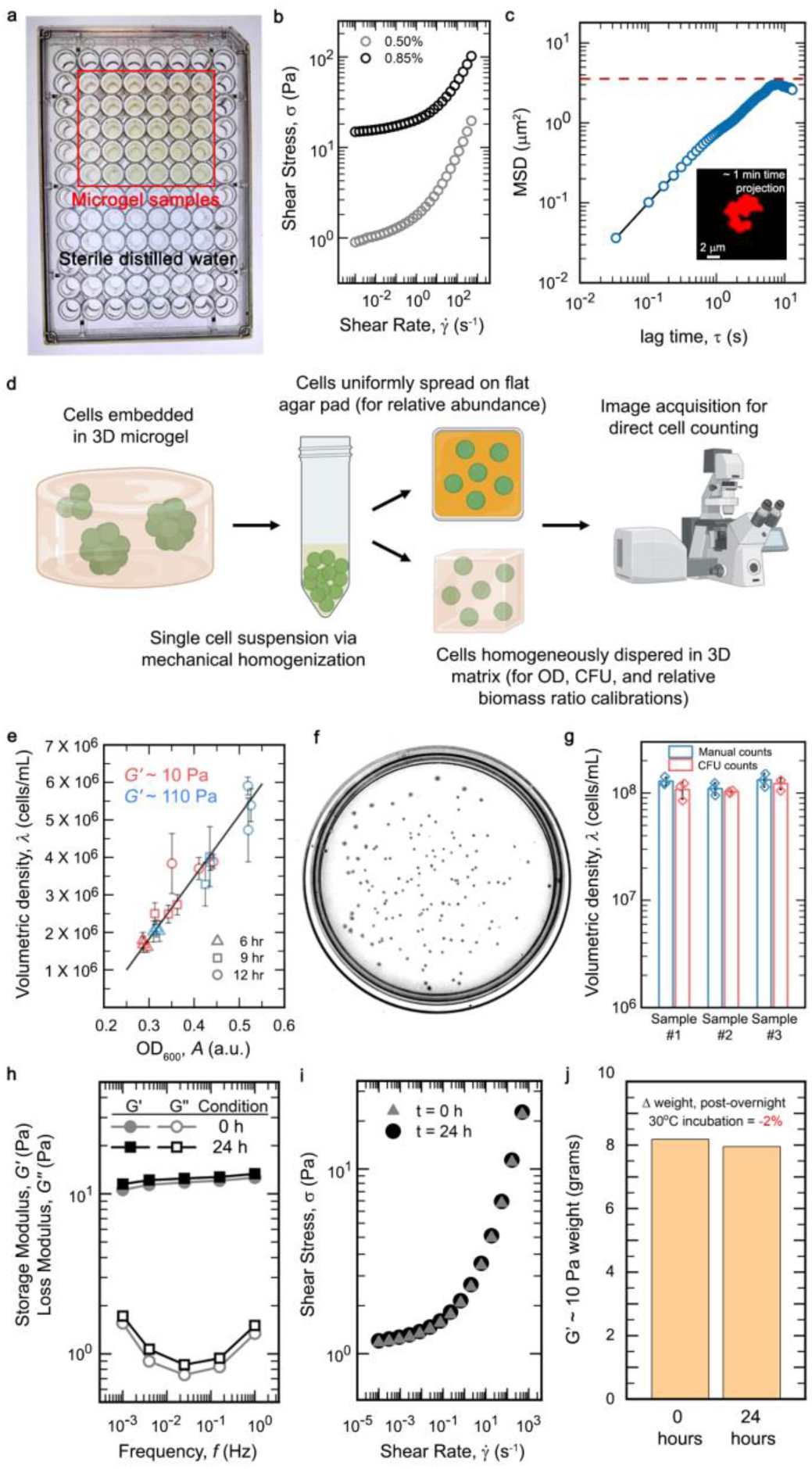
(a) Representation of the microgel-based culturing setup, wherein, bacteria-laden microgel samples are maintained in 96 well plates under humid conditions (by filling up surrounding wells with sterile distilled water) in 30°C to minimize evaporative losses. (b) Rheological measurements reveal that the 3D growth media behaves as a yield stress material – i.e., it undergoes reversible fluidization at high shear rates while behaving as a soft solid at low shear rates. (c) Representative particle tracking of a 200nm fluorescent tracer particle moving via thermal diffusion within the inter-particle pore spaces of the 3D microgel matrix (n = 55 individual particles were imaged for each gel type). The plateau value represents the cage size of the pore that constrains the overall particle displacement. (d) Schematic representation of sample processing for direct cell counting. Sub-panels of this illustration were prepared using Biorender (2024), BioRender.com/n92g221. (e) A comparison between absorbance measurements and volumetric density of cells shows a broadly linear relation for both the low confinement and high confinement matrices, indicating that the optical density-based absorbance measurements can be used as a reliable indicator of total bacterial biomass for the 3D growth media. Volumetric density is measured by directly counting the total number of cells present in a 3D volumetric image from confocal microscopy (n = 3 technical replicates for absorbance measurements per each time point, and n = 4 distinct fields of view for volumetric cell counting acquired from each individual replicate of the absorbance measurement samples). (f and g) We directly benchmark the microscopic 3D volumetric counting method against conventional colony forming units (CFU)-based estimation of cell numbers and show good agreement between both (n = 3 distinct fields of view for volumetric cell counting, and n = 3 technical replicates for the colony forming units assay plates). (h, i, and j) Culturing conditions do not cause appreciable changes to the rheological properties or bulk volume of the 3D microgel matrix, even for the low confinement system.

**Extended data Figure 3:**
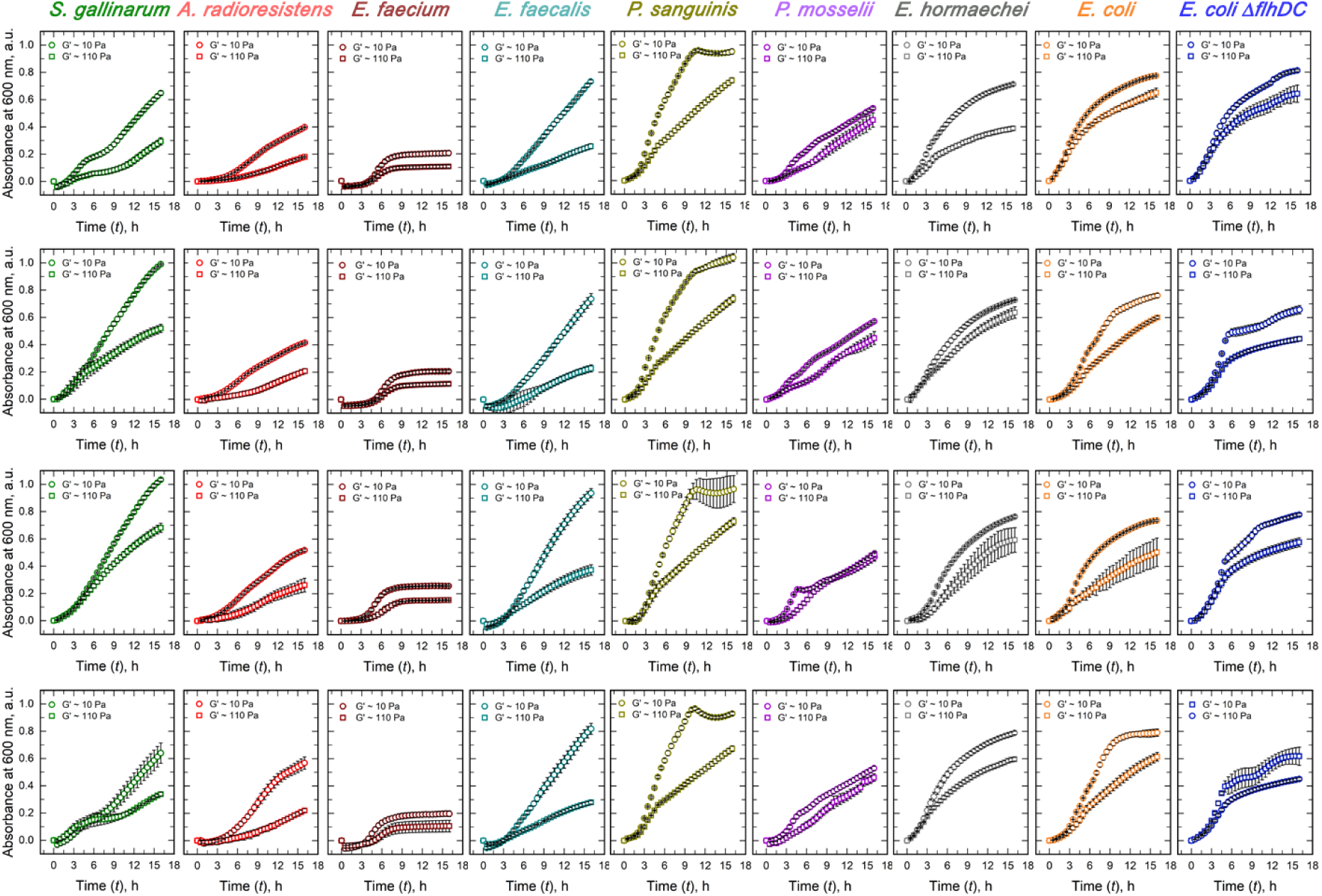
Representative biological replicates from growth curve assays in 3D microgel growth media for all bacterial strains used in this study (mean +/- s.d., for n = 2-4 technical replicates for each biological replicate).

**Extended data Figure 4:**
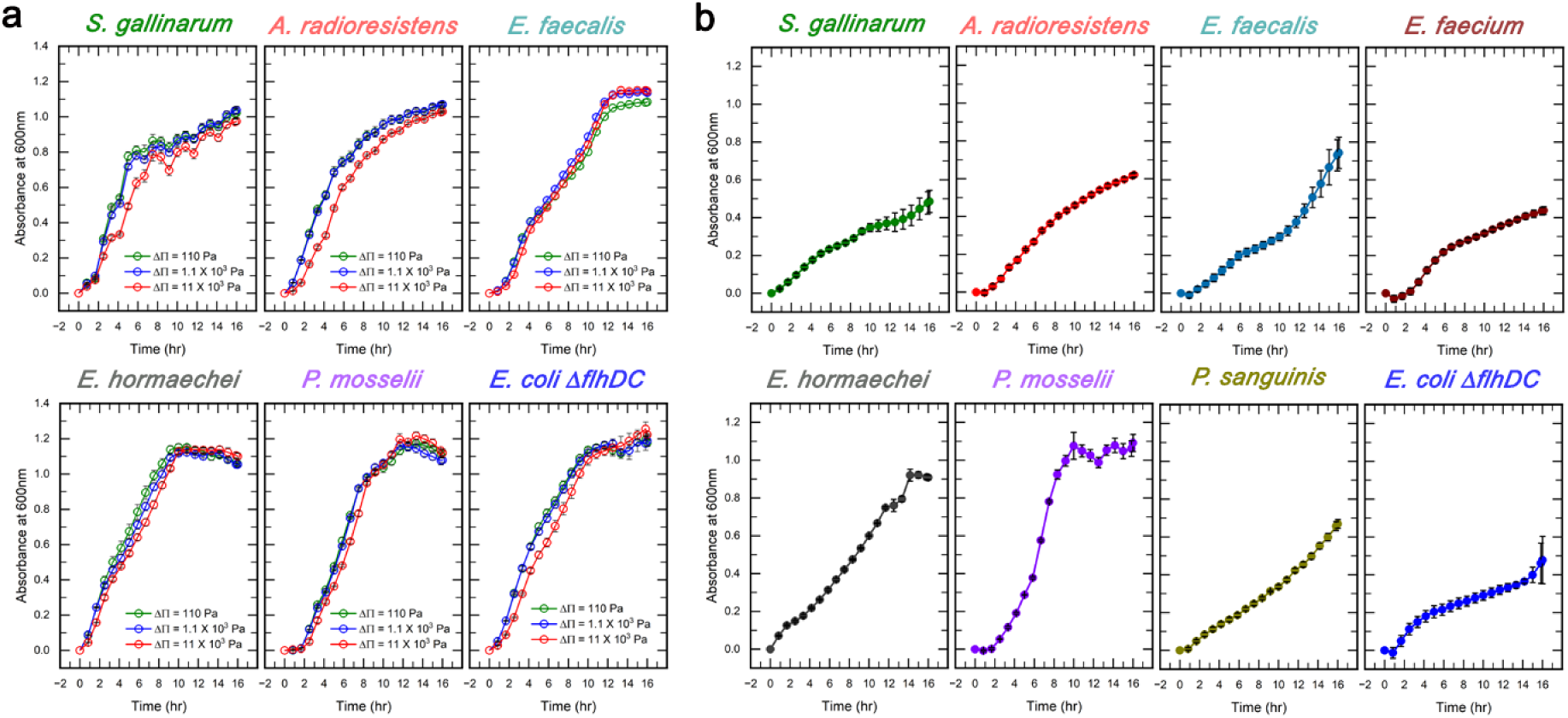
(a) Growth curves of three spherical and three rod-shaped bacterial strains in shaken liquid LB, with different degrees of osmotic pressure elevations achieved by adding the inert polymer PEG1000 to these cultures (mean +/- s.d., for n = 3 technical replicates). No significantly detrimental effect of increased osmotic pressure on bacterial growth is observed. (b) Non-shaken liquid LB cultures of all bacterial strains used in this study. Static culturing conditions in a liquid media do not promote effective microbial growth (mean +/- s.d., for n = 3 technical replicates).

**Extended data Figure 5:**
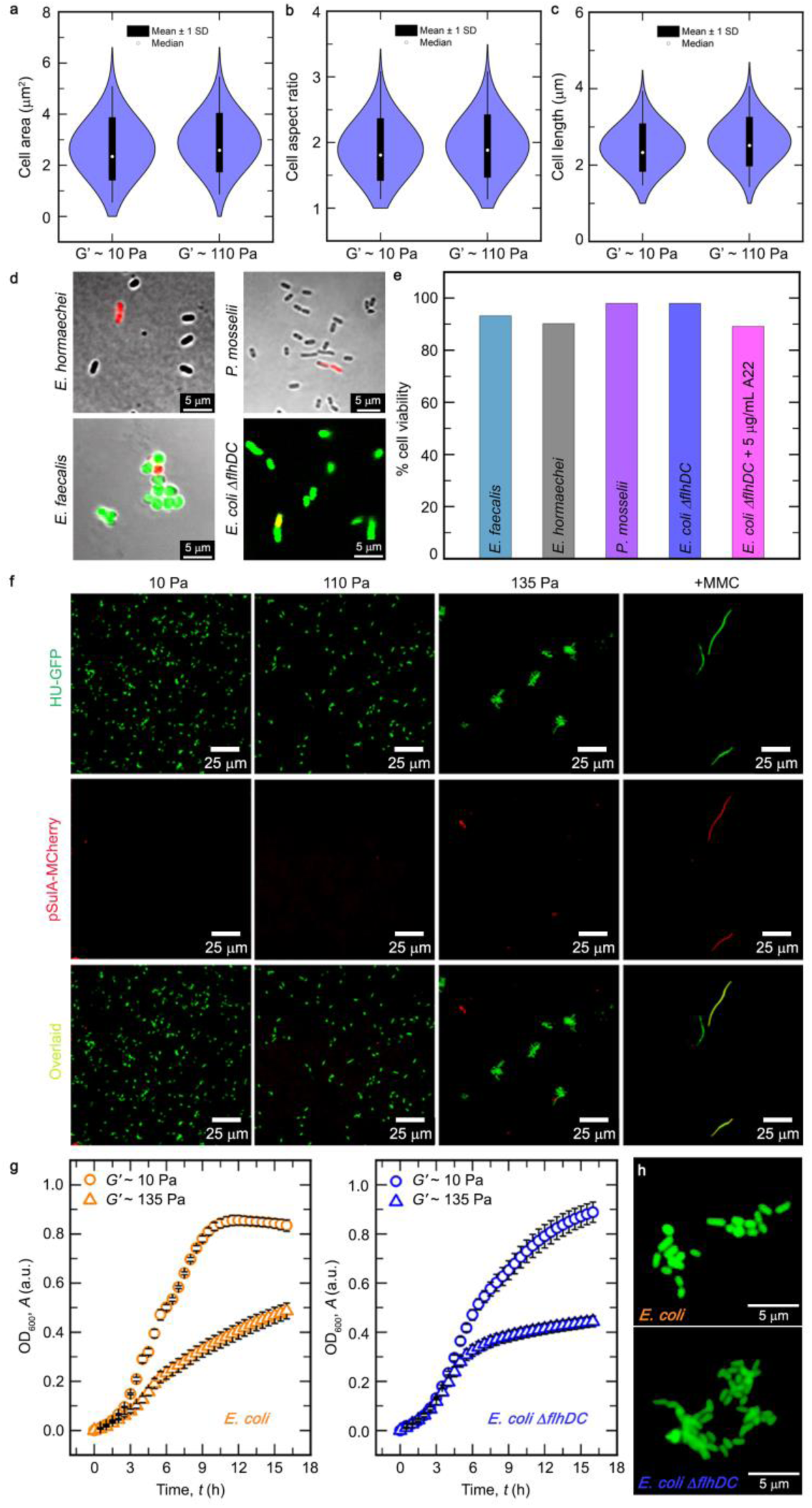
(a-c) The degree of physical confinement within jammed microgel growth media does not alter the cellular morphology of bacterial cells, as shown here for non-motile *E. coli* cells (n > 300 individual cells across both gel types represented as normal distribution, with boxes indicating the means, and bounds of the boxes indicating +/- s.d., and circles indicating the median). (d and e) Growth under 3D confinement does not detrimentally affect the viability of bacterial cells. Staining using propidium iodide (red) as a marker of membrane-compromised cells (presumed dead), combined with calcein-AM (green) for *E. faecalis* and constitutive GFP expression for *E. coli* show that following 24 hours of growth in a high confinement matrix, population-level cell viability remains high (∼=/> 90%) across several different microbial strains used in this study (n > 240 individual cells for all strain types). (f) Growth under confinement does not affect the genomic integrity of bacteria, as evidenced by the lack of DNA damage response reporter (SulA, tagged to MCherry) expression. The HU-GFP indicates the genome organisation, since HU is a DNA binding protein. Hence, the signal in the green fluorescence channel indicates the genomic organisation of the *E. coli* grown under confinement. The SulA-Mcherry indicates triggering of DNA damage responses, hence the signal in the red fluorescence channel marks out cells which show DNA damage response. Since we did not observe any noticeable levels of this signal from cells grown under physical confinement in our 3D microgel matrices, we went a step further and directly induced DNA damage to cells grown in liquid using the drug MMC, which has been included as a positive control. Representative micrographs from a single set of experiments. (g) Motility does not confer significantly greater growth benefits under confinement (mean +/- s.d., for n = 3 technical replicates). Using highly confined microgel growth media, we observe that both motile and non-motile *E. coli* perform similarly. (h) Here we show that under high degrees (*G’* ∼ 135 Pa microgels) of 3D confinement, both motile and non-motile *E. coli* form elongated colonies. Typically, if a matrix allows for mobility, a motile strain is expected to disperse evenly and undertake single cell-driven growth. However, once we restrict the bacteria under sufficiently high degrees of 3D confinement, we observe that even motile bacteria are now forced to adopt a more localized, colony-driven form of growth, as their spatial dispersal is significantly hindered. Representative micrographs from a single set of experiments.

**Extended data Figure 6:**
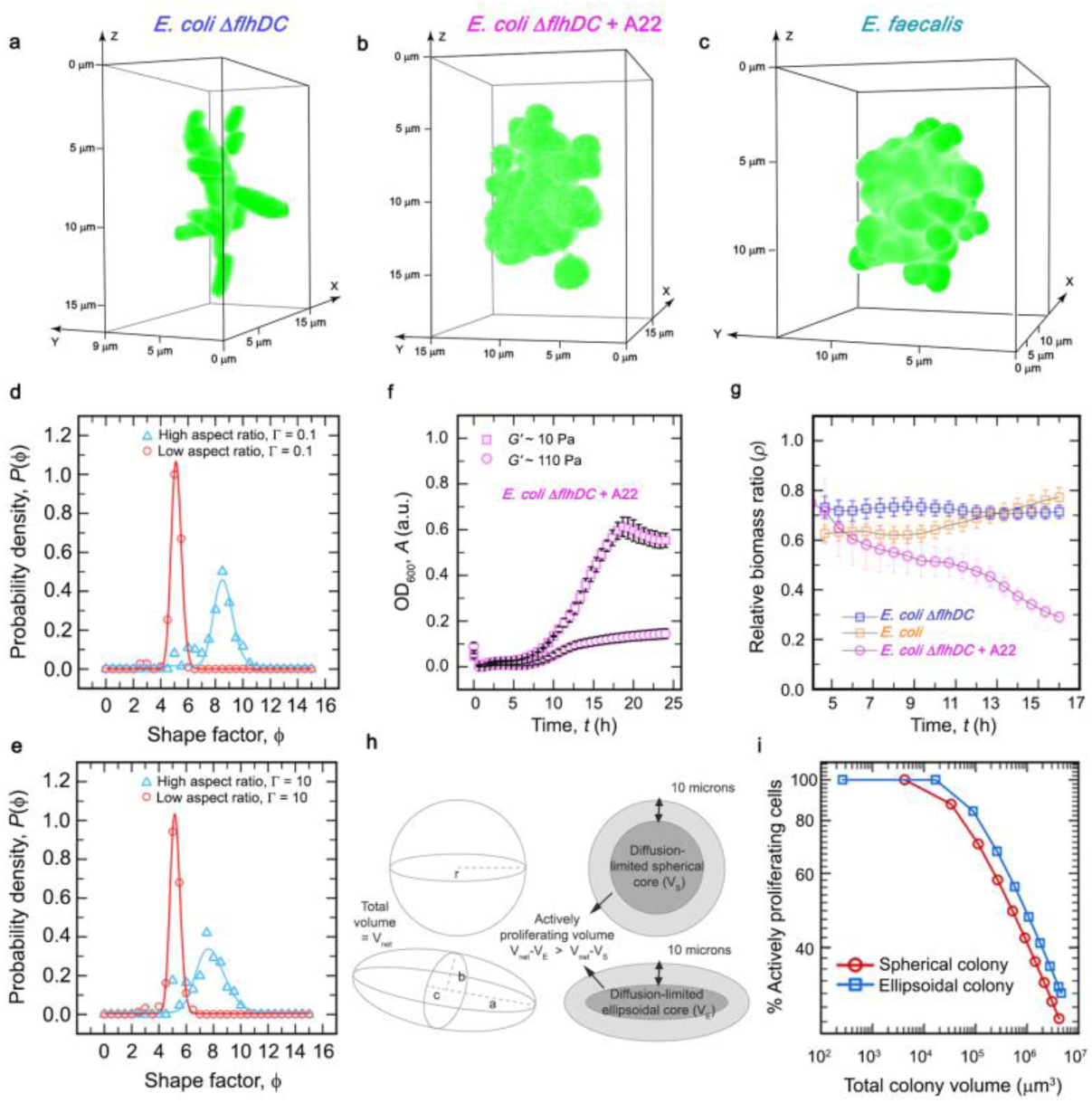
(a, b, and c) 3D reconstructions of colonies formed by rod-shaped *E. coli*, approximately spherical *E. coli*, and naturally spherical *E. faecalis* during growth in the high confinement matrix. Similar phenomena have been observed for at least three biological replicates; representative micrographs shown here for a single such colony for each strain. (d and e) Quantitative analysis of colony morphologies using the shape factor metric, generated using simulations with different conditions of environmental drag (*Γ*) on bacterial growth. In all cases, a clear distinction is visible between colonies formed by high vs. low aspect ratio bacteria, with the former organizing into colonies with a high shape factor value (more elongated), whereas, the latter organizes into more rounded colonies with a low shape factor value (n ≥ 200 snapshots, from 10 simulations). (f) Representative growth curves of *E. coli* treated with A22 under two different degrees of confinement, assayed for 24 hours of growth, considering the antibiotic-induced delay in proliferation (mean +/- s.d., for n = 3 technical replicates). (g) Relative biomass ratio (*ρ*) across 16 hours of growth under confinement for *E. coli*, non-motile *E. coli*, and non-motile *E. coli* treated with A22, showing how a shift in the bacterial shape towards approximately spherical morphology dramatically diminishes the relative growth success under increased confinement (mean +/- propagated error, which is calculated as described in the statistical analysis, with biological replicates n = 5 for rod-shaped motile and non-motile *E. coli*, as well as n = 4 for A22-treated spherical non-motile *E. coli*). (h) Schematic representations of numerical calculations considering either spherical colonies formed by low aspect ratio (=1) cells or ellipsoidal colonies formed by high aspect ratio (=4) cells. From previous work on *E. coli* confined in 3D porous media, we assume a maximum depth of ∼10 μm up to which nutrient diffusion allows active proliferation of cells, beyond which the colony’s core remains nutrient limited and hence metabolically inactive. (i) The proportion of actively proliferating cells in colonies of different sizes for either condition, showing that ellipsoidal colonies formed by high aspect ratio cells retain a significantly higher proportion of cells in the actively proliferating fraction, compared to spherical colonies formed by low aspect ratio cells, for colonies of comparable sizes.

**Extended data Figure 7:**
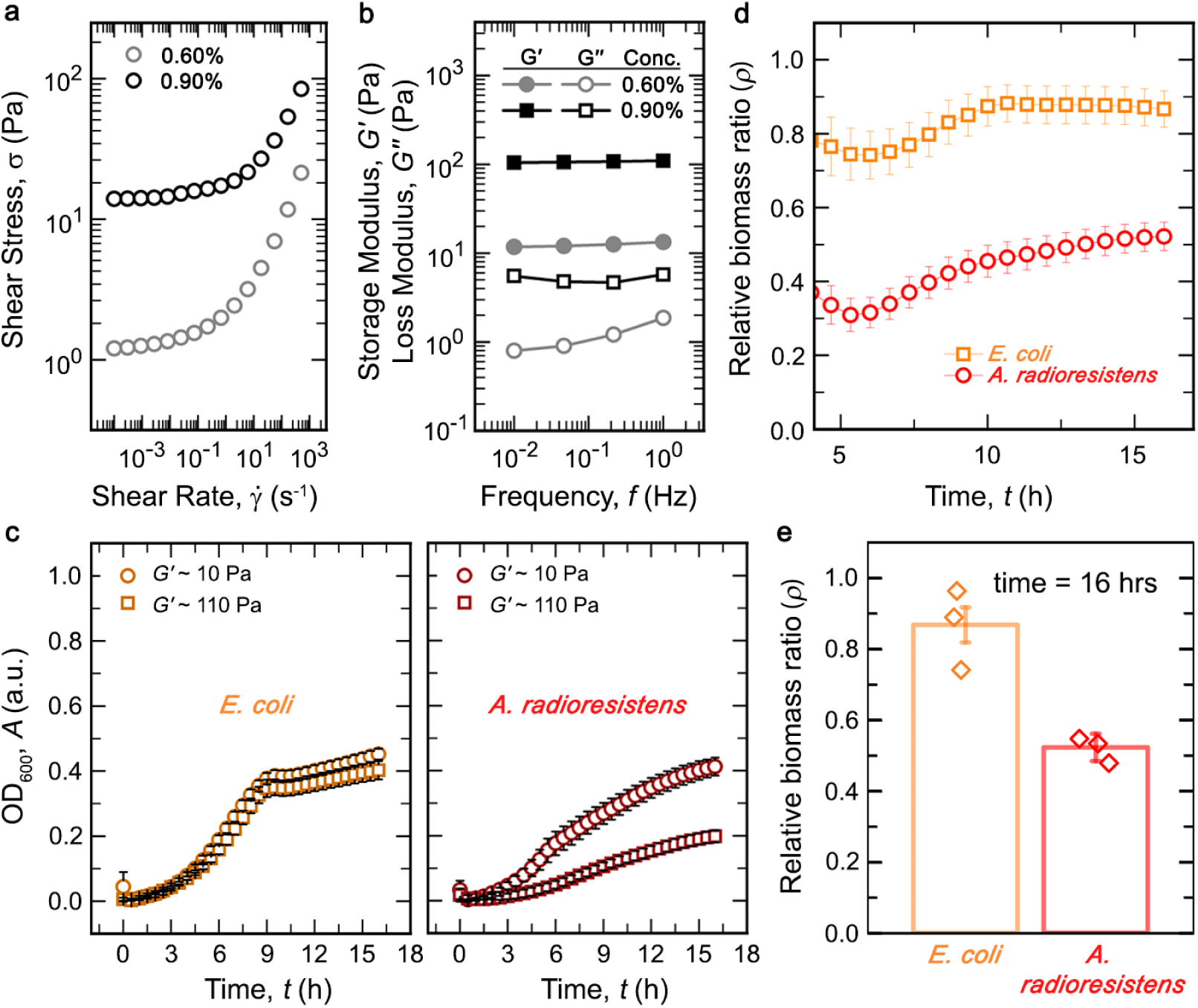
Characterizing bacterial growth dynamics in minimal media (M9)-based jammed microgel matrix. We replicate conditions of confinement similar to those imposed by the LB-based microgels using M9-based 3D growth matrix. (a and b) Rheological characterization of M9-based microgels reveals physical properties matching that of LB-based 3D growth media. (c) To test whether the trends exhibited by relative biomass ratios for low and high aspect bacteria are independent of nutrient composition, we assay the high aspect ratio *E. coli* and low aspect ratio *A. radioresistens* in minimal media (M9) supplemented with 0.2% glucose and 0.2% milk protein hydrolysate (mean +/- s.d., for n = 3 technical replicates). (d and e) Relative biomass ratio trends over 16 hours of growth for the rod-shaped *E. coli* and spherical *A. radioresistens* across two different degrees of confinement using M9-based jammed microgel growth matrix (mean +/- propagated error, which is calculated as described in the statistical analysis, with biological replicates n = 3 for rod-shaped motile *E. coli* and spherical *A. radioresistens*). The results recapitulate what has been observed using LB-based 3D growth matrix, with the high aspect ratio *E. coli* exhibiting a superior relative biomass ratio under a higher degree of confinement.

**Extended data Figure 8:**
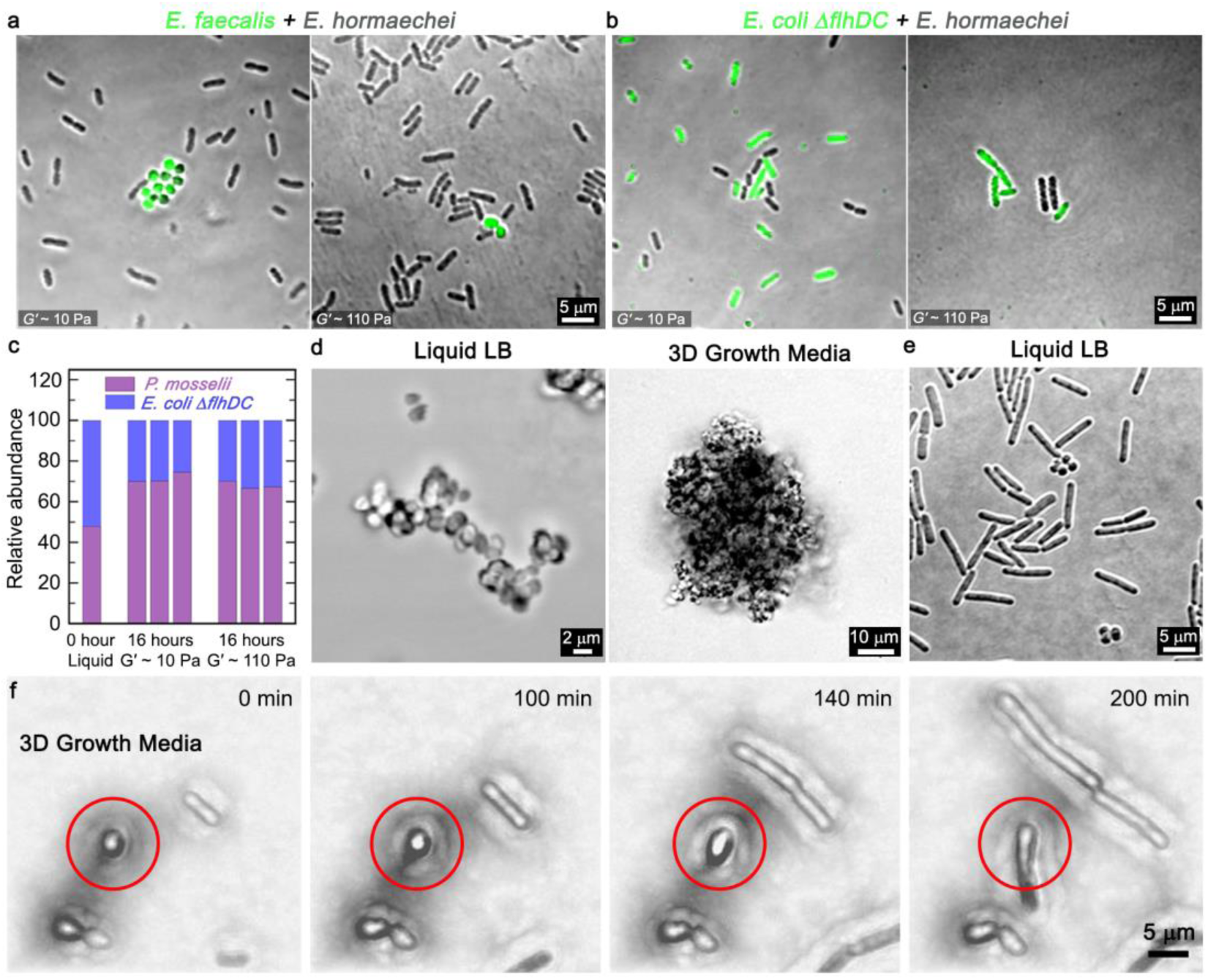
(a, b, and c) 3D co-culture models show that high aspect ratio cells (motile *E. hormaechei*) enjoy a distinct advantage over low aspect ratio cells (*E. faecalis*) upon an increase in physical confinement (for *E. hormaechei* vs *E. faecalis*, n = 2 biological replicates, each with >= 400 individual cells pooled from >= 10 unique micrographs for each gel type). By contrast, competitive cultures between two different rod-shaped species – *E. coli* and *E. hormaechei*, (for non-motile *E. coli* vs *E. hormaechei*, n = 2 biological replicates, each with >= 130 individual cells pooled from >= 10 unique micrographs for each gel type), or, *E. coli* and *P. mosselii* (n = 3 biological replicates) – does not show a significant enrichment of either cell type. (d) *E. faecalis*, which exhibits growth as elongated clusters in liquid LB (representative micrograph from a single experiment), instead shows colony growth in the form of spherical colonies in the 3D growth media under physical confinement (representative micrograph from at least three biological replicates). (e and f) Growth under confinement leads to dramatically altered outcomes at both the colony and single-cell levels. *P. sanguinis* shows a mixed spherical and rod-shaped morphology in liquid LB, but within microgel growth media, exhibits highly filamentous growth, resulting in elongated colonies (representative micrographs from at least two biological replicates).

**Extended data Table 1:**
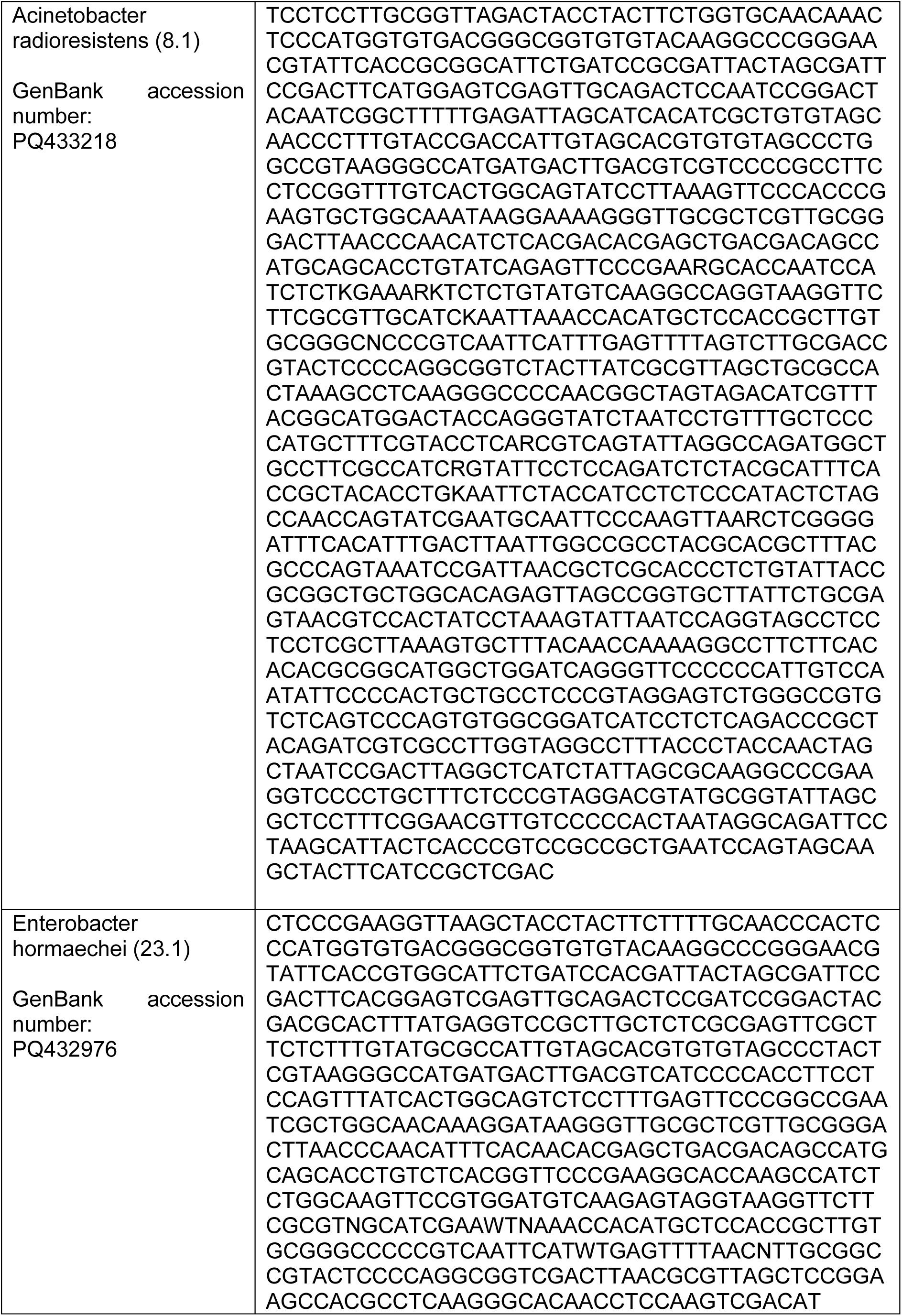
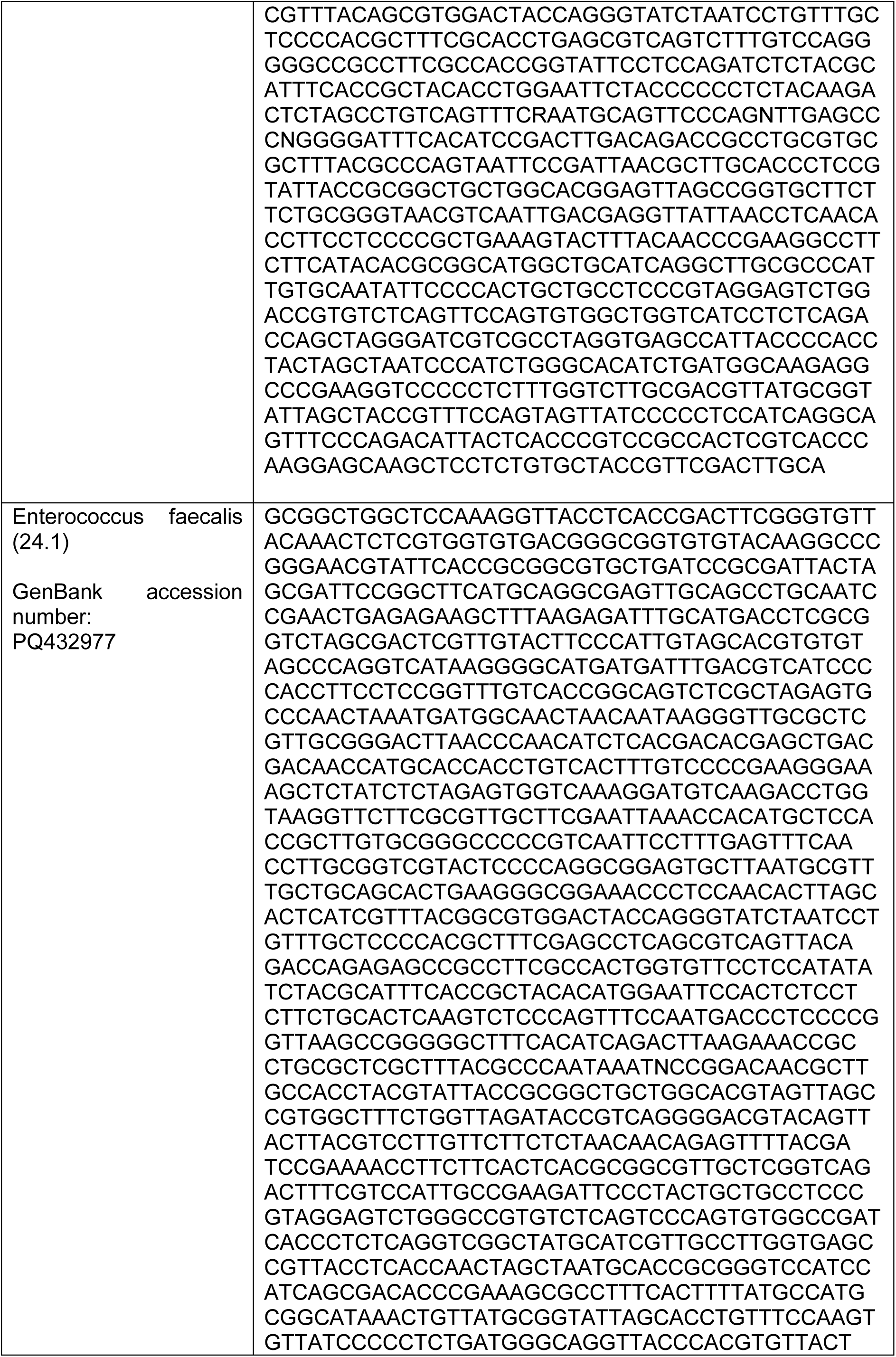
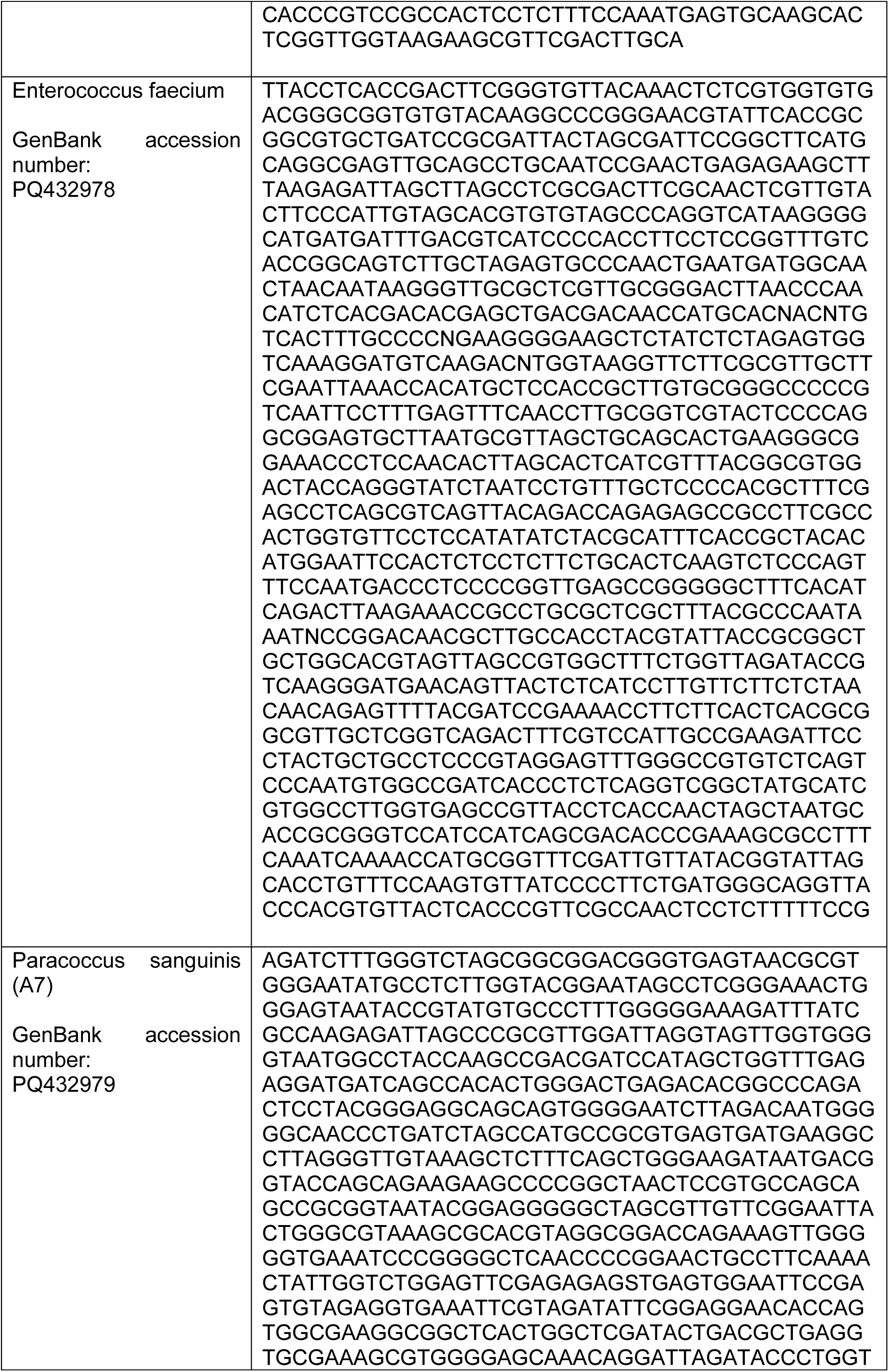
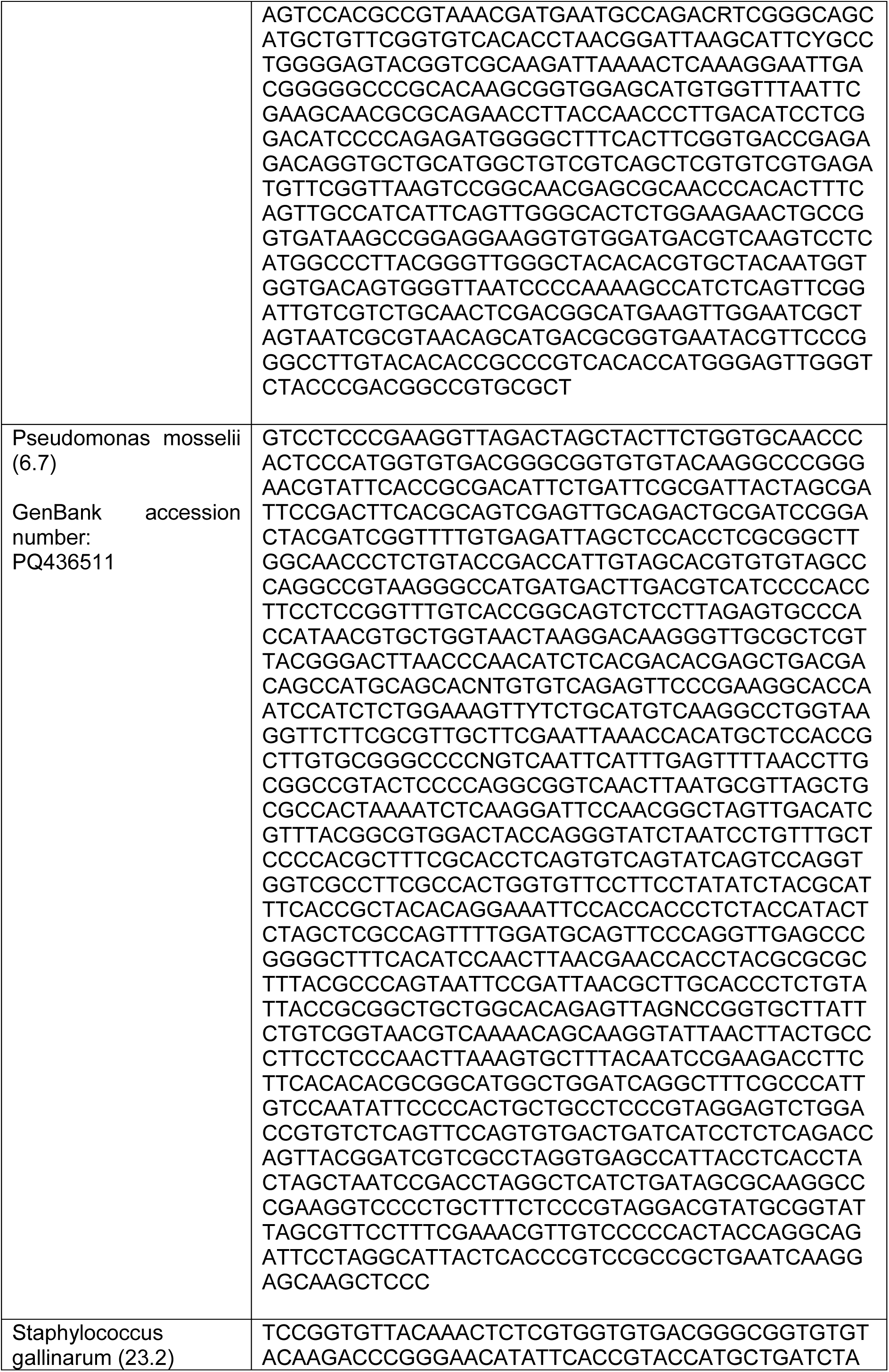
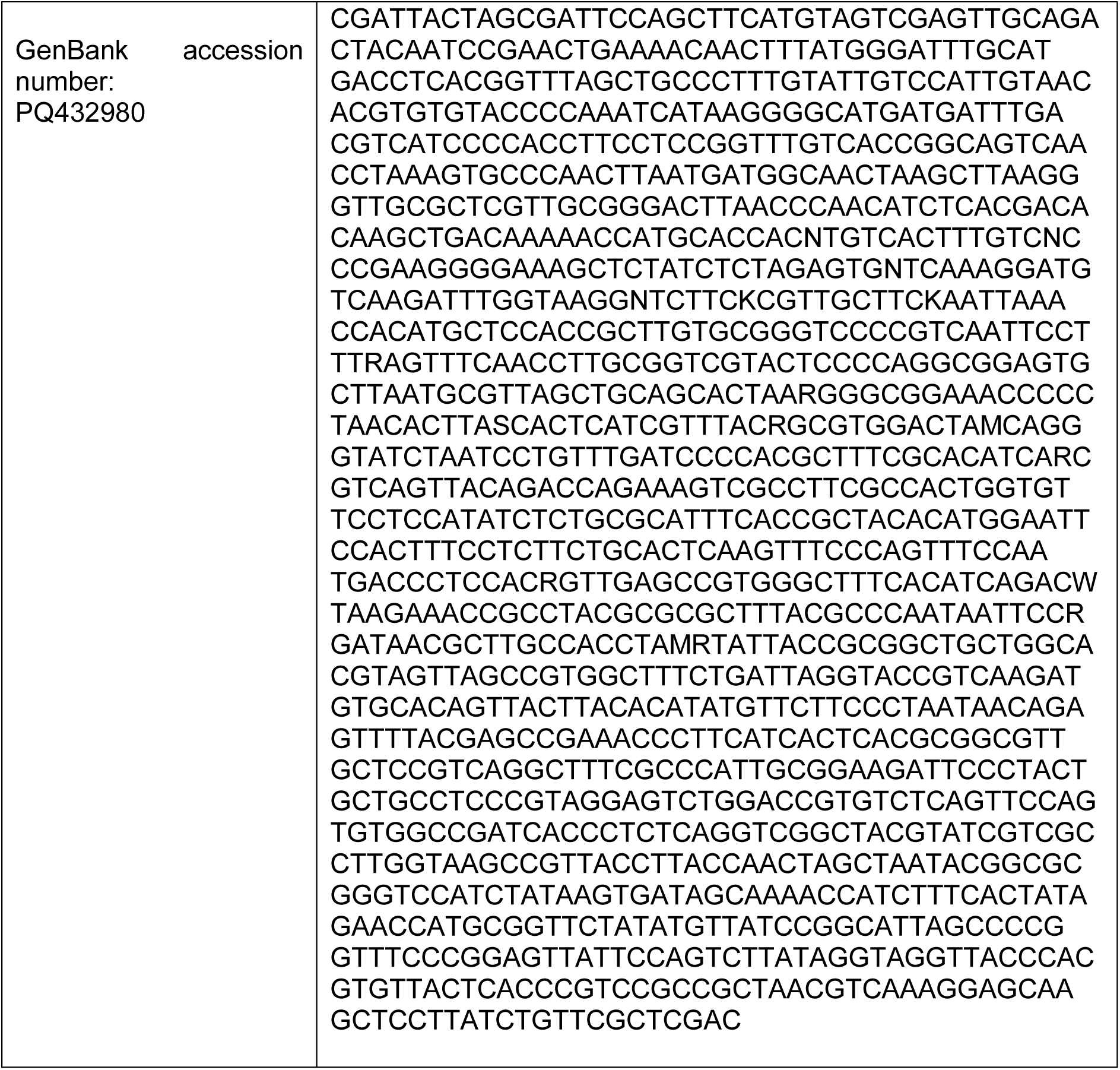
16S rRNA Sequencing of all beetle-derived microbial strains.

## Supplementary videos

**Supplementary video 1:** Simulations of bacterial colony growth in 3D.

**Supplementary video 2:** Microscopic imaging of bacterial colony growth inside 3D matrices.

**Supplementary video 3:** Time lapse imaging of *P. sanguinis* growth inside the 3D microgel matrices.

## References

1. Leff, J. W. et al. Consistent responses of soil microbial communities to elevated nutrient inputs in grasslands across the globe. Proc Natl Acad Sci U S A 112, 10967– 10972 (2015).

2. Estrela, S., Sanchez-Gorostiaga, A., Vila, J. C. C. & Sanchez, A. Nutrient dominance governs the assembly of microbial communities in mixed nutrient environments. Elife 10, (2021).

3. Keegstra, J. M., Carrara, F. & Stocker, R. The ecological roles of bacterial chemotaxis. Nature Reviews Microbiology 2022 20:8 20, 491–504 (2022).

4. Gonzalez, D. & Mavridou, D. A. I. Making the Best of Aggression: The Many Dimensions of Bacterial Toxin Regulation. Trends Microbiol 27, 897–905 (2019).

5. Blasche, S. et al. Metabolic cooperation and spatiotemporal niche partitioning in a kefir microbial community. Nature Microbiology 2021 6:2 6, 196–208 (2021).

6. Chang, C. Y., Bajić, D., Vila, J. C. C., Estrela, S. & Sanchez, A. Emergent coexistence in multispecies microbial communities. Science (1979) 381, 343–348 (2023).

7. Persat, A. et al. The Mechanical World of Bacteria. Cell 161, 988 (2015).

8. Datta, S. S., Steinberga, A. P. & Ismagilova, R. F. Polymers in the gut compress the colonic mucus hydrogel. Proc Natl Acad Sci U S A 113, 7041–7046 (2016).

9. Ribet, D. & Cossart, P. How bacterial pathogens colonize their hosts and invade deeper tissues. Microbes Infect 17, 173–183 (2015).

10. Gomez, S. et al. Substrate stiffness impacts early biofilm formation by modulating Pseudomonas aeruginosa twitching motility. Elife 12, (2023).

11. Tchoufag, J., Ghosh, P., Pogue, C. B., Nan, B. & Mandadapu, K. K. Mechanisms for bacterial gliding motility on soft substrates. Proc Natl Acad Sci U S A 116, 25087– 25096 (2019).

12. Flemming, H. C. et al. Biofilms: an emergent form of bacterial life. Nature Reviews Microbiology 2016 14:9 14, 563–575 (2016).

13. Schloss, P. D. & Handelsman, J. Toward a Census of Bacteria in Soil. PLoS Comput Biol 2, e92 (2006).

14. Fan, Y. & Pedersen, O. Gut microbiota in human metabolic health and disease. Nature Reviews Microbiology 2020 19:1 19, 55–71 (2020).

15. Nicholson, P. G. Soil Improvement and Ground Modification Methods. Soil Improvement and Ground Modification Methods (2015).

16. Lu, P., Takai, K., Weaver, V. M. & Werb, Z. Extracellular matrix degradation and remodeling in development and disease. Cold Spring Harb Perspect Biol 3, (2011).

17. Belkaid, Y. & Hand, T. W. Role of the microbiota in immunity and inflammation. Cell 157, 121–141 (2014).

18. Kreda, S. M., Davis, C. W. & Rose, M. C. CFTR, Mucins, and Mucus Obstruction in Cystic Fibrosis. Cold Spring Harb Perspect Med 2, a009589 (2012).

19. Shao, X. et al. Growth of bacteria in 3-d colonies. PLoS Comput Biol 13, e1005679 (2017).

20. Martinez-Calvo, A. et al. Morphological instability and roughening of growing 3D bacterial colonies. Proc Natl Acad Sci U S A 119, e2208019119 (2022).

21. Kannan, H. et al. Spatiotemporal development of growth and death zones in expanding bacterial colonies driven by emergent nutrient dynamics. bioRxiv 2023.08.27.554977 (2023) doi:10.1101/2023.08.27.554977.

22. Dell’Arciprete, D. et al. A growing bacterial colony in two dimensions as an active nematic. Nature Communications 2018 9:1 9, 1–9 (2018).

23. You, Z., Pearce, D. J. G., Sengupta, A. & Giomi, L. Mono-to Multilayer Transition in Growing Bacterial Colonies. Phys Rev Lett 123, 178001 (2019).

24. Bhattacharjee, T. & Datta, S. S. Bacterial hopping and trapping in porous media. Nature Communications 2019 10:1 10, 1–9 (2019).

25. Lai, S. K., Wang, Y. Y., Hida, K., Cone, R. & Hanes, J. Nanoparticles reveal that human cervicovaginal mucus is riddled with pores larger than viruses. Proc Natl Acad Sci U S A 107, 598–603 (2010).

26. Sardelli, L. et al. Towards bioinspired in vitro models of intestinal mucus. RSC Adv 9, 15887–15899 (2019).

27. Bhattacharjee, T. & Datta, S. S. Confinement and activity regulate bacterial motion in porous media. Soft Matter 15, 9920–9930 (2019).

28. Bhattacharjee, T. et al. Polyelectrolyte scaling laws for microgel yielding near jamming. Soft Matter 14, 1559–1570 (2018).

29. Bhattacharjee, T. et al. Liquid-like Solids Support Cells in 3D. ACS Biomater Sci Eng 2, 1787–1795 (2016).

30. Sreepadmanabh, M., Ganesh, M., Bhat, R. & Bhattacharjee, T. Jammed microgel growth medium prepared by flash-solidification of agarose for 3D cell culture and 3D bioprinting. Biomedical Materials 18, 045011 (2023).

31. Sreepadmanabh, M., Arun, A. B. & Bhattacharjee, T. Design approaches for 3D cell culture and 3D bioprinting platforms. Biophys Rev 5, (2024).

32. Sleator, R. D. & Hill, C. Bacterial osmoadaptation: the role of osmolytes in bacterial stress and virulence. FEMS Microbiol Rev 26, 49–71 (2002).

33. Wood, J. M. Bacterial responses to osmotic challenges. Journal of General Physiology 145, 381–388 (2015).

34. Osawa, M. & Erickson, H. P. Turgor pressure and possible constriction mechanisms in bacterial division. Front Microbiol 9, 331686 (2018).

35. Kamashev, D., Balandina, A. & Rouviere-Yaniv, J. The binding motif recognized by HU on both nicked and cruciform DNA. EMBO J 18, 5434–5444 (1999).

36. Burby, P. E. & Simmons, L. A. Regulation of cell division in bacteria by monitoring genome integrity and DNA replication status. J Bacteriol 202, (2020).

37. Rudge, T. J., Steiner, P. J., Phillips, A. & Haseloff, J. Computational Modeling of Synthetic Microbial Biofilms. ACS Synth Biol 1, 345–352 (2012).

38. Pinho, M. G., Kjos, M. & Veening, J. W. How to get (a)round: mechanisms controlling growth and division of coccoid bacteria. Nature Reviews Microbiology 2013 11:9 11, 601–614 (2013).

39. Bean, G. J. et al. A22 disrupts the bacterial actin cytoskeleton by directly binding and inducing a low-affinity state in MreB. Biochemistry 48, 4852–4857 (2009).

40. Khandige, S. et al. DamX controls reversible cell morphology switching in uropathogenic escherichia coli. mBio 7, 642–658 (2016).

41. Lee, Y. & Wang, C. Morphological Change and Decreasing Transfer Rate of Biofilm-Featured Listeria monocytogenes EGDe. J Food Prot 80, 368–375 (2017).

42. Espinosa, E. et al. l-Arabinose Induces the Formation of Viable Nonproliferating Spheroplasts in Vibrio cholerae. Appl Environ Microbiol 87, 1–15 (2021).

43. Khandige, S. et al. DamX Controls Reversible Cell Morphology Switching in Uropathogenic Escherichia coli. mBio 7, (2016).

44. Bockwoldt, J. A., Zimmermann, M., Tiso, T. & Blank, L. M. Complete Genome Sequence and Annotation of the Paracoccus pantotrophus Type Strain DSM 2944. Microbiol Resour Announc 9, (2020).

45. McGinnis, J. M. et al. Paracoccus sanguinis sp. nov., isolated from clinical specimens of New York state patients. Int J Syst Evol Microbiol 65, 1877–1882 (2015).

46. Nokhal, T. H. & Mayer, F. Structural analysis of four strains of Paracoccus denitrificans. Antonie Van Leeuwenhoek 45, 185–197 (1979).

47. Quinn, R. A. et al. Niche partitioning of a pathogenic microbiome driven by chemical gradients. Sci Adv 4, (2018).

48. Baran, R. et al. Exometabolite niche partitioning among sympatric soil bacteria. Nature Communications 2015 6:1 6, 1–9 (2015).

49. Brochet, S. et al. Niche partitioning facilitates coexistence of closely related gut bacteria. Elife 10, (2021).

50. Nair, R. R. et al. Bacterial predator-prey coevolution accelerates genome evolution and selects on virulence-associated prey defences. Nature Communications 2019 10:1 10, 1–10 (2019).

51. Wongkiew, S., Chaikaew, P., Takrattanasaran, N. & Khamkajorn, T. Evaluation of nutrient characteristics and bacterial community in agricultural soil groups for sustainable land management. Scientific Reports 2022 12:1 12, 1–13 (2022).

52. Battesti, A., Majdalani, N. & Gottesman, S. The RpoS-mediated general stress response in escherichia coli *. Annu Rev Microbiol 65, 189–213 (2011).

53. Benomar, S. et al. Nutritional stress induces exchange of cell material and energetic coupling between bacterial species. Nature Communications 2015 6:1 6, 1–10 (2015).

54. Peterson, S. B., Bertolli, S. K. & Mougous, J. D. Interbacterial antagonism: at the center of bacterial life. Curr Biol 30, R1203 (2020).

55. MacLean, R. C., Torres-Barceló, C. & Moxon, R. Evaluating evolutionary models of stress-induced mutagenesis in bacteria. Nature Reviews Genetics 2013 14:3 14, 221–227 (2013).

56. Foster, P. L. Stress-Induced Mutagenesis in Bacteria. Crit Rev Biochem Mol Biol 42, 373 (2007).

57. Tran, T. D., Ali, M. A., Lee, D., Félix, M. A. & Luallen, R. J. Bacterial filamentation as a mechanism for cell-to-cell spread within an animal host. Nature Communications 2022 13:1 13, 1–11 (2022).

58. Smith, W. P. J. et al. Cell morphology drives spatial patterning in microbial communities. Proc Natl Acad Sci U S A 114, E280–E286 (2017).

59. Monier, J. M. & Lindow, S. E. Differential survival of solitary and aggregated bacterial cells promotes aggregate formation on leaf surfaces. Proc Natl Acad Sci U S A 100, 15977–15982 (2003).

60. Rutherford, S. T. & Bassler, B. L. Bacterial Quorum Sensing: Its Role in Virulence and Possibilities for Its Control. Cold Spring Harb Perspect Med 2, (2012).

61. Li, S., Liu, S. Y., Chan, S. Y. & Chua, S. L. Biofilm matrix cloaks bacterial quorum sensing chemoattractants from predator detection. ISME J 16, 1388 (2022).

62. Matz, C. et al. Marine Biofilm Bacteria Evade Eukaryotic Predation by Targeted Chemical Defense. PLoS One 3, 2744 (2008).

63. Gray, D. A. et al. Extreme slow growth as alternative strategy to survive deep starvation in bacteria. Nature Communications 2019 10:1 10, 1–12 (2019).

64. Crocker, J. C. & Grier, D. G. Methods of Digital Video Microscopy for Colloidal Studies. J Colloid Interface Sci 179, 298–310 (1996).

65. Naik, T., Sharda, M., Lakshminarayanan, C. P., Virbhadra, K. & Pandit, A. High-quality single amplicon sequencing method for illumina MiSeq platform using pool of ‘N’ (0-10) spacer-linked target specific primers without PhiX spike-in. BMC Genomics 24, (2023).

66. Callahan, B. J. et al. DADA2: High-resolution sample inference from Illumina amplicon data. Nature Methods 2016 13:7 13, 581–583 (2016).

